# Rapid in-EPON CLEM for everyone: Combining fast and efficient labeling of self-labeling enzyme tags with EM-resistant Janelia Fluor dyes

**DOI:** 10.1101/2023.11.04.565612

**Authors:** Rico Franzkoch, Sabrina Wilkening, Viktoria Liss, Michael Holtmannspötter, Rainer Kurre, Olympia E. Psathaki, Michael Hensel

## Abstract

Correlative light and electron microscopy (CLEM) allows to link light microscopy (LM) of living cells to ultrastructural analyses by electron microscopy (EM). Pre-embedding CLEM often suffers from inaccurate correlation between the LM and EM modalities due to chemical and physical distortions. Post-embedding CLEM enables precise registration of fluorescent structures directly on thin resin sections. However, in-resin CLEM techniques require fluorescent markers withstanding EM sample preparation. Most fluorescent proteins lose their fluorescence during EM sample preparation. Synthetic dyes present an alternative as their photostability and brightness exceed those of fluorescent proteins. Together with self-labeling enzymes (SLE) as protein tags, these fluorophores can be used to precisely label cellular structures of interest. By applying SLE labelling for post-embedding CLEM, we compared Janelia Fluor dyes and TMR to identify most suitable fluorophores. Epithelial cells expressing HaloTag fusion proteins were stained with various ligand-conjugated dyes, and fluorescence preservation was quantified after conventional room temperature sample preparation with embedding in EPON. The results obtained show that only the red dyes TMR, JF549, JFX549 and JFX554 retain their fluorescence in resin, with JFX549 and JFX554 yielding best signal intensity and signal-to-background ratio during in-resin super-resolution microscopy. Since all red dyes possess an oxygen atom within their xanthene structure, our results indicate that this might be a crucial feature making them more tolerant to sample preparation for electron microscopy. Our work reports a rapid in-resin CLEM approach that combines fast and efficient labeling of SLE tags with EM-compatible fluorophores, and serve as benchmarks for experimental planning and future engineering of fluorophores for CLEM.

## Introduction

Transmission electron microscopy (TEM) with its near atomic spatial resolution has been used successfully for decades to unravel the ultrastructural architecture of different cellular components (Winey et al., 2014). The chemicals used in sample preparation for EM interact with and thereby stain a variety of cellular content enabling the visualization of nearly all cellular components at the same time (Bozzola & Kuo, 2014). This results in a unique reference space. The localization of specific proteins on TEM sections is possible for instance via immunogold labeling, but needs specifically adapted protocols, suitable antibodies, a very careful interpretation of the results, and is limited through the penetration depth of the antibodies (Schwarz & Humbel, 2007; Stierhof & Schwarz, 1989; Tokuyasu, 1980). In light microscopy (LM) cellular structures or proteins can be directly labeled by fluorescent proteins (FP) or dyes with high efficiency and the dynamics inside living cells can be easily monitored (Liss et al., 2015). Due to the comparably low resolution of LM and the lack of ultrastructural context precise and unambiguous identification of the structures underlying the fluorescence signal remains challenging. The combination of both imaging modalities as correlative light and electron microscopy (CLEM) is able to overcome the individual limitations and presents a powerful tool gaining more and more recognition in cellular biology during the recent years (de Boer et al., 2015; Ganeva & Kukulski, 2020; Krieger et al., 2014). Depending on the biological question to be addressed, several CLEM approaches were developed. These can be distinguished in workflows conducting the LM prior to embedding (pre-embedding), after the embedding step (post-embedding), or employing a combination of both. During pre-embedding CLEM, the sample is either imaged in living or aldehyde-fixed state. The former allows for visualization of cellular dynamics and does not impede the intensity of the fluorescence signal. Routinely following LM is EM sample preparation consisting of post-fixation with osmium tetroxide, dehydration, and final embedding in resin (Bozzola, 2014; Krieger et al., 2014). Such steps are known to induce artifacts including shrinkage and extraction of cellular material (McDonald & Auer, 2006). Such artifacts can severely compromise correlation, especially since Z resolution in LM is ca. 500 nm, thus about 10-times lower than for ultrathin sections of typical thickness of 50-70 nm. To overcome these limitations, post-embedding LM in combination with high-pressure freezing (HPF) and freeze substitution (FS) is an emerging alternative. For this, LM and EM modalities are registered on the same section, resulting in very high accuracy of correlation (Buerger et al., 2021; Kukulski et al., 2011). Furthermore, ultrastructural preservation is highly improved due to HPF and FS (McDonald & Auer, 2006). Drawbacks of such techniques are the need for sophisticated, expensive equipment, and use of methacrylate resins (e.g. Lowicryl HM20) which also limit contrast and have inferior properties in sectioning compared to epoxy resins. Also, the increased complexity of workflows demands highly experienced personnel (Bykov et al., 2016; Tanida et al., 2020). Certain staining and fixation agents like osmium tetroxide have to be completely omitted, or can only be used in low concentrations such as 0.1% uranyl acetate, since these chemicals have a strong impact on the fluorescence (Fu et al., 2020; Heiligenstein & Lucas, 2022; Kukulski et al., 2011).

A recent approach to simplify post-embedding CLEM is to use fluorescent proteins which can, to some degree tolerate the quenching properties of conventional EM sample preparation protocols including osmium tetroxide staining and dehydration at room temperature. For this, special variants of the Eos-FP like mEosEM were engineered or standard fluorescent proteins like mWasabi, mKate2 or mScarlet-H were tested for their resistance to TEM sample preparation (Fu et al., 2020; Paez-Segala et al., 2015; Peng et al., 2022; Tanida et al., 2020). However, many of these proteins seem to retain only comparably weak fluorescence signals and low signal to background ratio after EPON embedding, and require protocols with reduced osmium tetroxide concentration (Peng et al., 2022).

An alternative to fluorescent proteins are fluorophores such as tetramethylrhodamine (TMR) that can be coupled to ligands that are specifically, covalently, and irreversibly bound by self-labeling enzyme tags (SLE) such as HaloTag, SNAP-tag or CLIP-tag (Liss et al., 2015). This system allows for rapid and precise labeling of tagged cellular structures, and is compatible with super-resolution microscopy (SRM). Regarding the use conventional TEM sample preparation protocols with EPON embedding, systematic analysis of performance of available fluorophore-conjugated SLE ligands is missing. Only few publications employed dyes in such workflows, suggesting that they might be promising alternatives to fluorescent proteins (Müller et al., 2017; Sanada et al., 2022). Müller et al. showed that fluorescent signals of insulin granules labelled with TMR-Star and 505-Star are visible after EPON embedding, and also after HPF and FS. However, in this conventional sample preparation protocol only low concentration (0.1%) of osmium tetroxide in combination with uranyl acetate was used for 30 min, which might not be sufficient to preserve all ultrastructural details. Additionally, very high concentrations (6 µM) of dyes and long incubation times (overnight) were employed, making this a more costly and time-consuming addition to the protocol. Sanada et al. and Tanida et al. employed higher osmium concentrations, but used cell-impermeable dyes such as DyLight 549 or HiLyte 555 which require cell permeabilization, and thereby possibly diminishes ultrastructural quality.

As we see great potential in a rapid in-resin CLEM approach that combines fast and efficient labeling of SLE tags with EM-resistant dyes, and the limited or partly contradictory published data, we set out to systematically evaluate the performance of various fluorescent SLE ligand conjugated dyes after TEM sample preparation with EPON embedding. We aimed for short labeling times to minimize additional workload in EM sample preparation, and use of low concentrations of dyes. For this, we especially focused on Janelia Fluor dyes since these were specifically engineered to exhibit the highest brightness and photostability outperforming other current dyes (Grimm et al., 2020).

## Results

### Rapid and simple conventional sample preparation for in-resin CLEM

In this study, we evaluated the fluorescence retainment of TMR and various Janelia Fluor dyes conjugated to the HaloTag ligand (HTL) after labeling intracellular proteins of interest tagged with the HaloTag and conventional EM sample preparation with EPON embedding for TEM. Janelia Fluor dyes were specifically engineered to exhibit the highest brightness and photostability outperforming other current dyes (Grimm, Muthusamy, et al., 2017; Grimm et al., 2020; Grimm et al., 2021). For comparison we chose HTL-TMR as standard available in our lab. The dyes tested, namely the green fluorescent JF479, the red fluorescent JF549, JFX549, JFX554 and JF585, and the far-red fluorescent JF646, JFX646 and JFX650, are listed in Table 4 with their properties. In **Fig. S 1**, the chemical structures and the origin of rhodamine-derived Janelia Fluor dyes can be found (from now on the term dye will be used to describe the combination of a dye with its HTL-ligand if not stated otherwise).

In an initial examination of the dyes, we checked the intracellular fluorescence only. The epithelial cell line HeLa permanently transfected with Tom20-HaloTag was routinely used for labeling experiments throughout this paper. The optimal concentration and labeling duration for each dye depend on many different conditions, like cell type, protein tagged, SLE used, ligand and dye itself. Previous studies used comparable dyes for LM applications at concentrations between 10 and 500 nM, and stained for 10 – 30 min (Beatty, 2019; Broadbent et al., 2023; Grimm, Brown, et al., 2017; Grimm et al., 2020; Joshua et al., 2022; Ricker et al., 2022), in case up to 1 µM (Binns et al., 2020; Braselmann et al., 2018), to label proteins conjugated to HaloTag. We used a concentration of 100 nM for 30 min for all dyes to keep conditions comparable. These labeling conditions were previously reported by our lab for HTL-TMR to provide a sufficient fluorescence signal in LM (Liss et al., 2015) and should serve as a comparative value to find well working dyes. HeLa Tom20-HaloTag cells were stained with 100 nM of each dye for 30 min before fixation with 3% PFA for 15 min. Subsequent imaging by confocal laser-scanning microscopy (CLSM) showed fluorescence signals matching the distribution of mitochondria and being clearly distinct from the background (**Fig. S 2**). Only JF479 and JF585 presented an exception with a low fluorescence signal barely sufficient to identify the stained mitochondria. Also noteworthy was the increased fluorescence signal of JFX646 compared to JF646.

Aiming for a CLEM approach that can easily be used in any laboratory without additional equipment, a standard EM embedding protocol with slight adaptations was used (**Fig. 1**). After labeling HeLa Tom20-HaloTag cells with various dyes in the respective concentration, cells were either first fixed with 0.2% GA and 3% PFA for locating ROIs in LM if necessary or directly subjected to EM preparation. Subsequent EM sample preparation was comprised of fixation with 2.5% GA for 1 h, post-fixation with osmium tetroxide and potassium hexacyanoferrate, dehydration in a graded ethanol series, incubation in anhydrous acetone, infiltration with and polymerization of EPON. To minimize quenching of the fluorescence by osmium tetroxide, its incubation time was reduced to 30 min, during post-fixation instead of 1 h as described in standard embedding protocols (Bozzola, 2014; Krieger et al., 2014). We checked the membrane preservation and contrast after this sample preparation protocol and found well preserved ultrastructure (**Fig. S 2**). Different cell organelles such as mitochondria, nucleus, lysosomes, Golgi and also membrane contact sites are well visible even after a shortened osmium incubation (**Fig. S 2**). Finally, sections of 250 nm or 100 nm were cut and viewed in widefield LM followed by contrasting with 3% uranyl acetate for 30 min and 2% lead citrate for 20 min. The images acquired by TEM were finally overlaid and correlated with the fluorescence images of the sections.

**Fig. 1.**
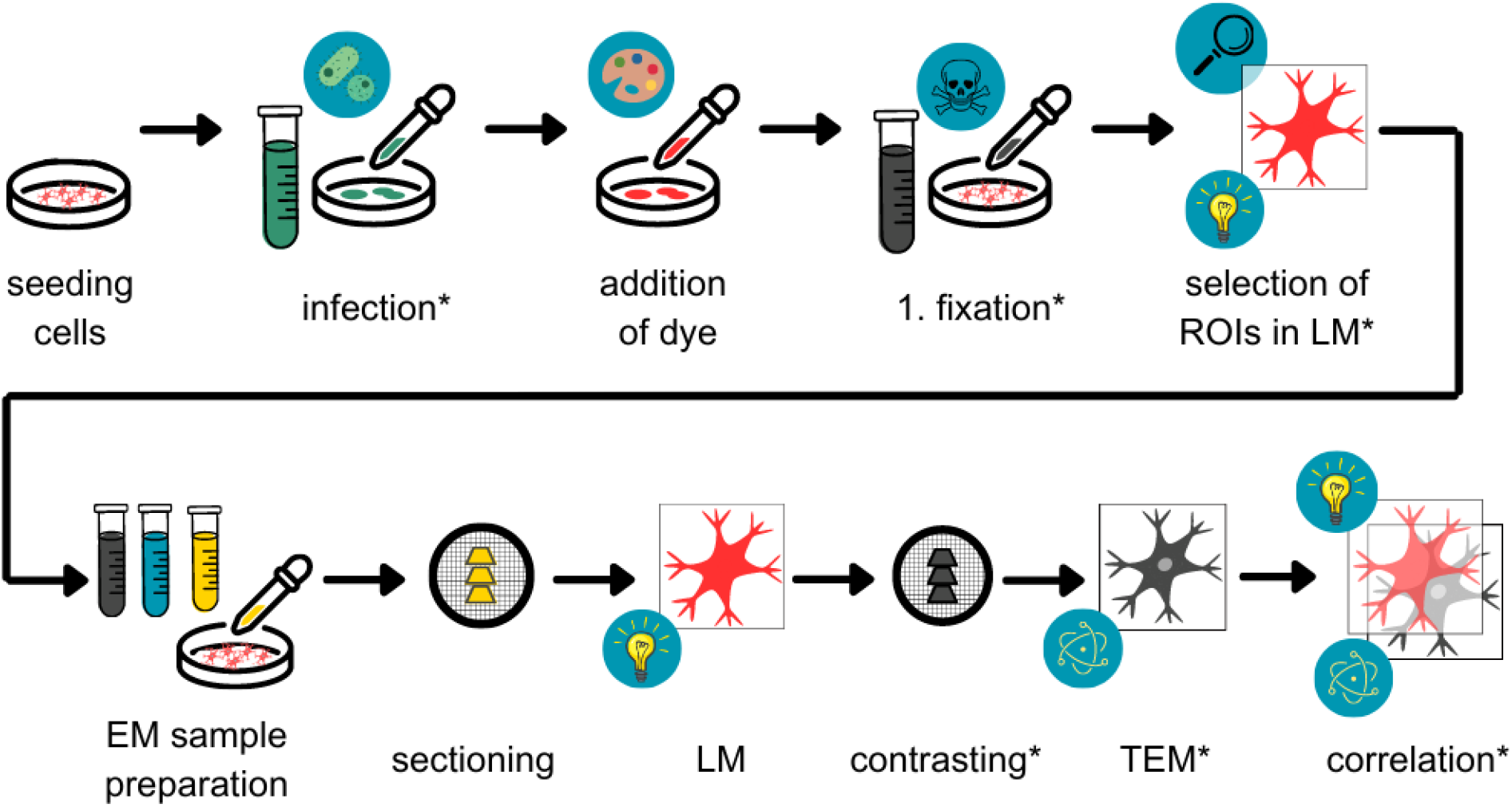
Workflow of sample preparation for CLEM. HeLa cells were optionally infected and stained before a first fixation with glutaraldehyde (GA) and paraformaldehyde (PFA). Cells of interest were selected by light microscopy (LM). Subsequently, the sample was prepared for electron microscopy (EM) by further fixation with GA, post-fixation with osmium tetroxide, dehydration in ethanol, and embedding in EPON. Sections were generated and viewed by widefield LM, followed by contrasting and imaging by transmission electron microscopy (TEM). Finally, the LM and EM modalities were overlaid and correlated. *Steps marked by an asterisk are omitted for quantitative comparison of various dyes.

### Certain Janelia Fluors retained fluorescence within thin EPON sections after conventional sample preparation

In a post-embedding CLEM approach evaluation of fluorescence retainment of dyes should be ideally done within the final sample. Therefore, we assessed the retained fluorescence directly on EPON sections. For that, HeLa Tom20-HaloTag cells were stained with the respective dye and prepared for EM as described in **Fig. 1**. Sections of 250 nm thickness were placed on coverslips or 50 mesh grids and analyzed by LM. Since pre-tests could not detect any fluorescence in embedded cells stained with 100 nM JF479 and the green dye already displayed a low fluorescence signal in first LM (**Fig. S 3**), JF479 was directly tested at a concentration of 1 µM and a staining duration of 2 h rather than 30 min to enhance the fluorescence signal of JF479 by counteracting the poor cell permeability of the dye. Yet, the increased concentration and the prolonged staining did not result in a fluorescence signal in the embedded sample, although the fluorescence signal was strongly enhanced before sample preparation. To exploit all possibilities, the LED power was additionally increased to 100% and exposure time was set to 1 s, compared to 30% and 500 ms, respectively, as used for the other dyes. Nevertheless, no fluorescence signals were observed for EPON sections of Tom20-JF479 (**Fig. 2**A). Increasing the concentration up to 10 µM, the upper reference value recommended by the supplier, still did not result in a detectable fluorescence signal in EPON sections (**Fig. 2**A).

**Fig. 2.**
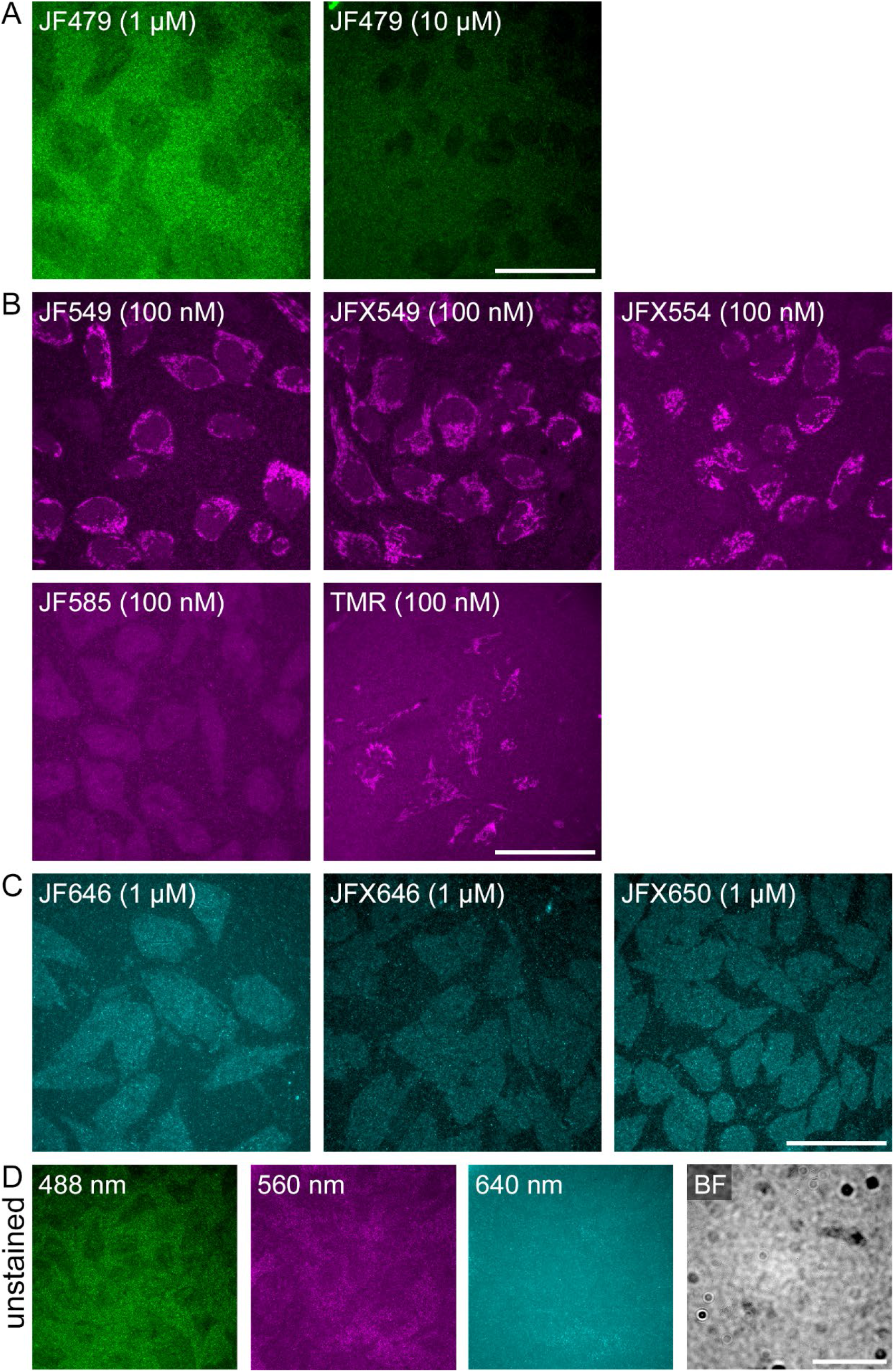
The red fluorescent dyes JF549, JFX549, JFX554, and TMR retain fluorescence after embedding in EM resin. HeLa cells expressing Tom20-HaloTag were stained with the indicated HTL conjugates with fluorescence emission in the green (**A**), red (**B**), or far-red (**C**) spectra at the indicated concentrations for 30 min. Cells were fixed with glutaraldehyde and osmium tetroxide, and embedded in EPON resin at 60 °C for 48 h. Next, the samples were sectioned into 250 nm sections, which were placed on a glass coverslip for imaging by widefield microscopy using an Olympus LSM FV3000 NLO. Analysis of the acquired images revealed loss of fluorescence for the green JF479 in both concentrations, for the fluorogenic red dye JF585, as well as for all tested far-red dyes, while fluorescence signals were detected for JF549, JFX549, JFX554, and TMR. **D**) An unstained control is shown that was processes identical to samples in **A**)-**C**), and imaged in the green, red, far-red, or brightfield (BF) channel. Scale bars: 50 µm.

For the red dyes JF549, JFX549, JFX554, and TMR, on the other hand, a fluorescence signal corresponding to a mitochondrial-like distribution was clearly detectable (**Fig. 2**B). The fluorogenic dye JF585 did not show any fluorescence at all and was comparable to the unstained control (**Fig. 2**B). Because JF585 already exhibited low fluorescence signals in LM, it was also examined at an increased concentration of 1 µM. Still, no fluorescence was observed on sections (**Fig. S 4**).

The results of JF646, JFX646, and JFX650 were similar to those obtained for JF479. Since pre-tests did not show any fluorescence on sections for 100 nM of ligand, concentration was increased to1 µM. Still no fluorescence was observed despite a longer exposure time of 1 s and higher LED power of 100% (**Fig. 2**C).

Apart from the fluorescence signal of the dyes, an uneven, speckle-like background signal was observed in the embedded sample. This background signal differed from the background signal within the cells (**Fig. 2**A-D). The background within the green channel was brighter than the background in the red channel, while no difference was observed in the far-red channel (**Fig. 2**).

### Quantification of fluorescence retainment

For the quantification of fluorescence preservation, the initial fluorescence intensity before EM sample preparation is compared to the signal after sample preparation, thus providing a value for the suitability of the dye. However, in post-embedding CLEM the fluorescence finally retained in resin sections is the most important value. Also, it is not possible to compare the fluorescence signal of a whole cell in liquid buffer with an ultrathin cell section in EPON due to several reasons, like different Z, different medium, usage of different LM settings and systems before and after EM sample preparation. In the past, some labs used an osmium tetroxide resistance assay (Fu et al., 2020; Peng et al., 2022; Tanida et al., 2020), however the impairment of the dye does not stop after the osmium step, but continues during additional steps of sample preparation. We thus only quantified fluorescence signals of dyes in 250 nm thin EPON sections, from at least three independent experiments and at least 100 cells per experiment, and compared them to the signal intensities of TMR. For this 250 nm thin sections were directly placed on a glass coverslip to facilitate LM. Registration of entire sections was conducted using widefield LM. Images were then analyzed with ImageJ following the algorithm described in Material and Methods.

Due to our aforementioned observations only the dyes TMR, JF549, JFX549 and JFX554 were used for quantification. Our tests revealed that JFX554 and JFX549 were the best performing dyes, retaining nearly 20% more fluorescence in EPON sections compared to TMR (**Fig. 3**A). Interestingly, we were also able to detect a significant difference between JF549 and the modified version JFX549 (**Fig. 3**A). Also note that TMR and JF549 did not show any fluorescence signals in 3 out of 6 and 3 out of 5 individual experiments, respectively, while retaining their fluorescence in the remaining experiments (**Fig. 3**). It seems that these two dyes are more susceptible during EM sample preparation than other dyes due to their structure and properties (**Fig. 3**). Due to the high background signal, the signal-to-background (S/B) ratio was computed for the four dyes. The S/B of JFX549 was slightly better than the S/B of JFX554 and of JF549, yet not significantly. Comparison of the S/B of these dyes with the S/B of TMR, which was the lowest, revealed a statistic significance at p < 0.001 (**Fig. 3**B).

**Fig. 3.**
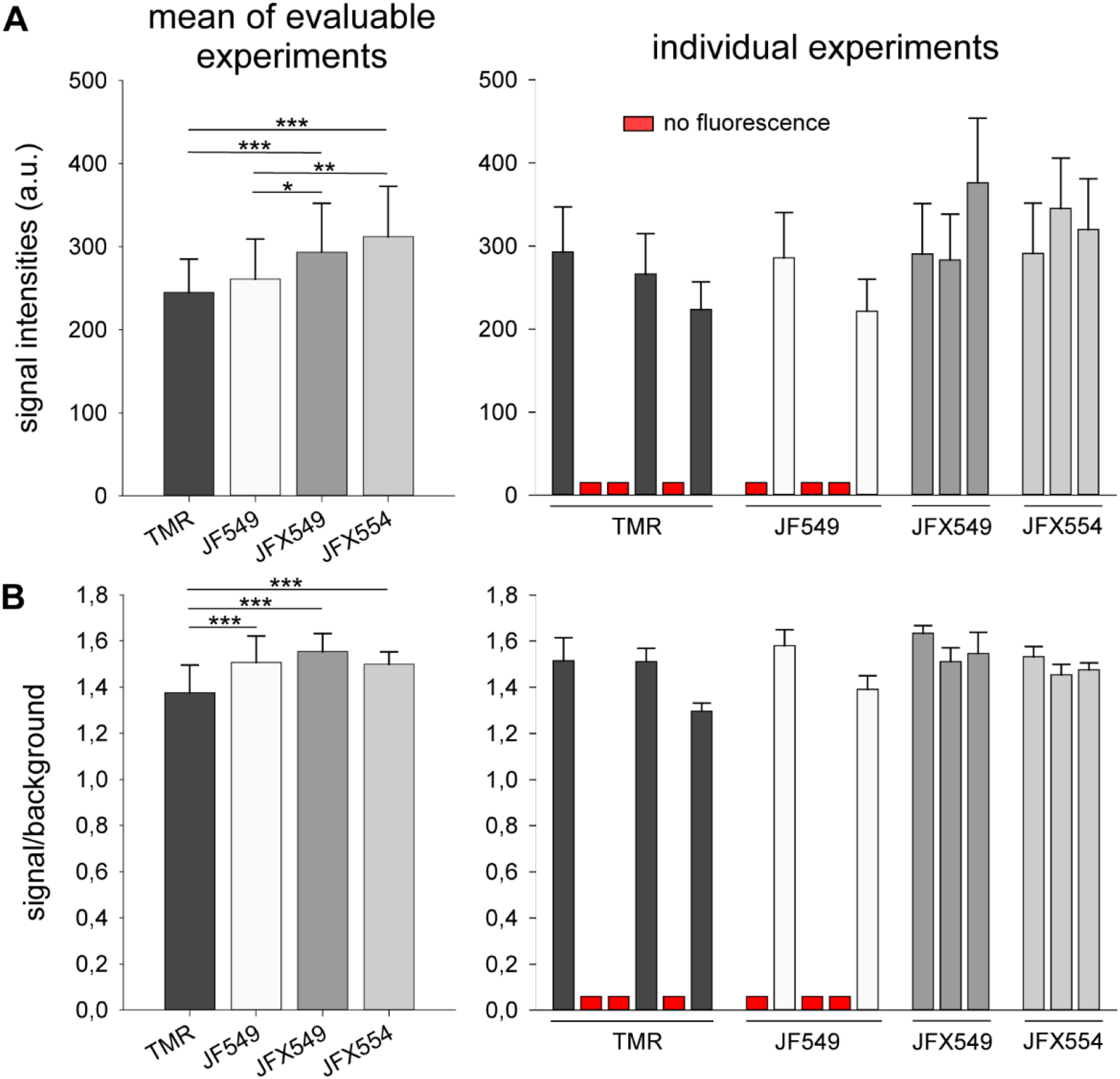
Comparison of fluorescence intensities and signal-to-background ratios of TMR, JF549, JFX549, and JFX554 in resin. The average signal intensity and standard deviations were ascertained by the algorithm for the selection and the background of each acquired image. **A**) Mean signal intensities of all evaluated images. The standard deviation was computed by averaging the standard deviation determined by the algorithm for the signal intensity of the selection and is indicated by the whiskers. ii) displays the same as i) but subdivided into the individual experiments. **B**) Signal-to-background ratios (S/B) was computed by dividing the average signal intensity of the selection by the average signal intensity of the background for each image. Results of individual experiments were shown in the right panel, and indicate that labeling with HTL-TMR or HTL-JF549 frequently failed, while JFX549 or JFX554 labelled in all cases. Only experiments resulting in detectable fluorescence were included in the calculation of means. Statistical analyses were performed by one-way ANOVA test and the Bonferroni t-test. Significances are indicated as follows: *, p < 0.05; **, p < 0.01; ***, p < 0.001.

To test if increased dye concentrations confer stronger fluorescence in resin, concentrations of 10 nM, 100 nM, or 1 µM for TMR and JFX554 were analyzed. Both dyes did not retain their fluorescence at 10 nM, which matches our observations of noticeable lower fluorescence signals at 10 nM in LM (data not shown). When comparing 100 nM to 1 µM TMR, a slight increase was found both in the fluorescence signal intensity and S/B ratio. JFX554, on the other hand, neither showed a statistically significant increase in the fluorescence signal, nor in the S/B ratio. Instead, the fluorescence signal rather decreased slightly (**Fig. S 5**).

In conclusion, at a concentration of 100 nM and 30 min of labeling, JFX549 and JFX554 were the best-performing dyes under our conditions, not only displaying the brightest fluorescence signal in EPON sections, but also providing consistent and reproducible results.

### In-resin CLEM

Based on the finding that JFX549 and JFX554 exhibited the highest in-resin fluorescence signals, we next examined their performance in representative CLEM workflows. HeLa Tom20-HaloTag cells were stained with 100 nM JFX554 for 30 min, EPON embedded, and sectioned into 100 nm and 250 nm thin sections. In contrast to the experimental procedure for quantitative analysis of the dye, now the resin sections were subsequently transferred to a formvar-coated 50 mesh grid, which was then placed between two glass coverslips with a drop of slightly alkaline PBS (pH 8.4) to improve fluorescence (Ader & Kukulski, 2017; Xiong et al., 2014). ROI of the sections were registered by LM, and after gold fiducial addition and contrasting imaged by TEM. Both semi-thin (250 nm) and ultra-thin (100 nm) sections displayed well detectable fluorescence signals in LM (**Fig. 4**ABCD). Nevertheless, a punctuate background, most likely resulting from the formvar coating and EPON embedding, was clearly visible and might complicate detection of smaller, punctuate structures (**Fig. 4**AC, yellow arrowheads). However, a precise correlation with standard TEM images and even tomography data was still very well possible, revealing single mitochondria in TEM positive for the fluorescence signal (**Fig. 4**Bi, Bii, Di, Dii). Depending on the sectioning angle and thickness of the section even differentiation between the mitochondrial matrix and the inner and outer membrane system was possible (**Fig. 4**D, Di, Dii). To overcome the speckled background on the sections we tried to place the sections on grids with a carbon film as done by many others for methacrylate sections to reduce background (Kukulski et al., 2011). Unexpectedly, this resulted in a grave degradation of the ultrastructure of the sample (**Fig. S 6A, B**). This effect seemed to be independent of the embedded biological material, since free EPON parts on the sections also looked degraded (**Fig. S 6C**) and the carbon film without any section on it was intact (**Fig. S 6D**). To test whether different parameters during LM induce this degradation we placed sections from the same sample onto formvar coated grids and subjected them to different steps of the LM workflow while also testing different pH of the buffer. No parameter degraded the ultrastructure on formvar coated grids, hinting into the direction that actually carbon coated grids and EPON sections might be incompatible (**Fig. S 6E – Ev**). LM of the liquid components of the used EPON resin does also not display the speckled background indicating that the final polymerization might be the crucial issue (data not shown). Furthermore, also EPON resin from other suppliers (Roth, Science Services) always gave comparable background (data not shown). To clarify that the aforementioned red dyes are indeed suitable for in-resin CLEM the CLEM workflow was conducted with TMR, JFX549 and in comparison, JFX554. For this experiment, no widefield LM but CLSM was used for the detection of the fluorescence signal, underlining that one is not restricted to a single LM technique. For all three dyes sufficient fluorescence signal was retained within 250 nm sections, allowing precise correlation in the TEM (**Fig. S 7**). One of the most convenient characteristics of LM is the possibility to differently label several structures at once and thereby visualizing their organization within a biological sample. Since the green and far-red Janelia Fluor dyes did not work in our setup while labeling mitochondria we set out to at least test another far-red dye, known to have worked in other CLEM approaches (Tanida et al., 2023), for its suitability in our in-resin CLEM workflow, namely Alexa Fluor 647. Alexa dyes are very prominent dyes in LM due to their brightness and bleaching properties. A great downside of those dyes however is their cell impermeability. To still visualize this dye inside cells we used Alexa Fluor 647 coupled to BSA and fed this mix to HeLa Tom20-HaloTag cells which were additionally stained with JFX554. Indeed, this approach allowed us to conduct dual-color in-resin CLEM clearly visualizing JFX554-positive mitochondria and Alexa Fluor 647-positive vesicles (**Fig. S 8**). Nevertheless, it is still unclear whether this dye is comparably resistant as e.g. JFX554 in conventional sample preparation, or if the high local concentration of the dye incorporated into vesicles prevented quenching of fluorescence signals during sample embedding.

**Fig. 4.**
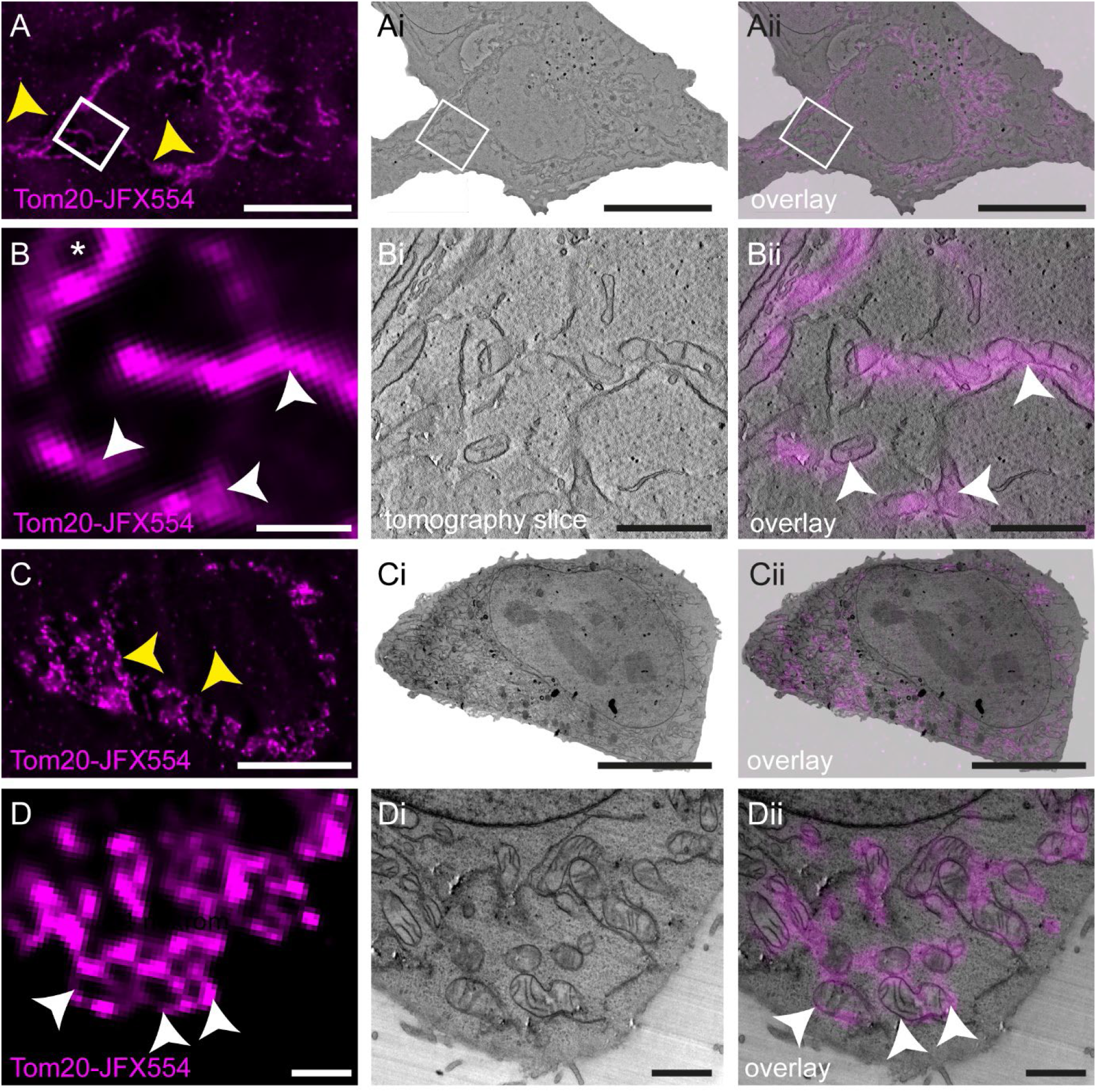
In-resin CLEM and tomography on 250 nm or 100 nm thin sections. HeLa cells stably expressing Tom20-HaloTag were stained with 100 nM JFX554 for 30 min and processed for in-resin CLEM. Sections of 250 nm (**A, B**) and 100 nm (**C, D**) thickness were prepared. After LM imaging (**A, C**) TEM tomography was conducted (**Ai, Bi,** see Movie 1 for tomogram), or ultra-thin sections were imaged using a standard TEM setup (**Ci, Di**). In both cases, fluorescence was retained after sample preparation and allow precise correlation (**Aii, Bii, Cii, Dii**). Scale bars: A, Ai, Aii, C, Ci, Cii: 10 µm; B, Bi, Bii, D, Di, Dii: 1 µm.

### Rhodamine dyes represent versatile markers for identification of different cellular structures during in-resin CLEM

To underline the versatility and general applicability of the here described in-resin CLEM workflow, different cellular structures were stained with JFX554 or TMR and visualized via in-resin CLEM (**Fig. 5**G). LM on 250 nm thin sections of HeLa cells expressing a Golgi-HaloTag marker revealed different Golgi stacks in a cell closely correlating with the underlying ultrastructure (**Fig. 5**A, Ai). TEM tomography of an exemplary region clearly depicted the membranes of the Golgi apparatus underlining the good contrast and preservation of the sample (Movie 2). In a HEK cell line expressing Connexin36 (Cx36) tagged with a SNAP-tag, extensive membrane whorls were detected in LM (**Fig. 5**B, C). Correlation with the underlying ultrastructure rapidly allowed identification of these structures and suggested them to be originating from the endoplasmic reticulum (**Fig. 5**Bi, Ci), which is in line with recently published data (Tetenborg et al., 2022).

**Fig. 5.**
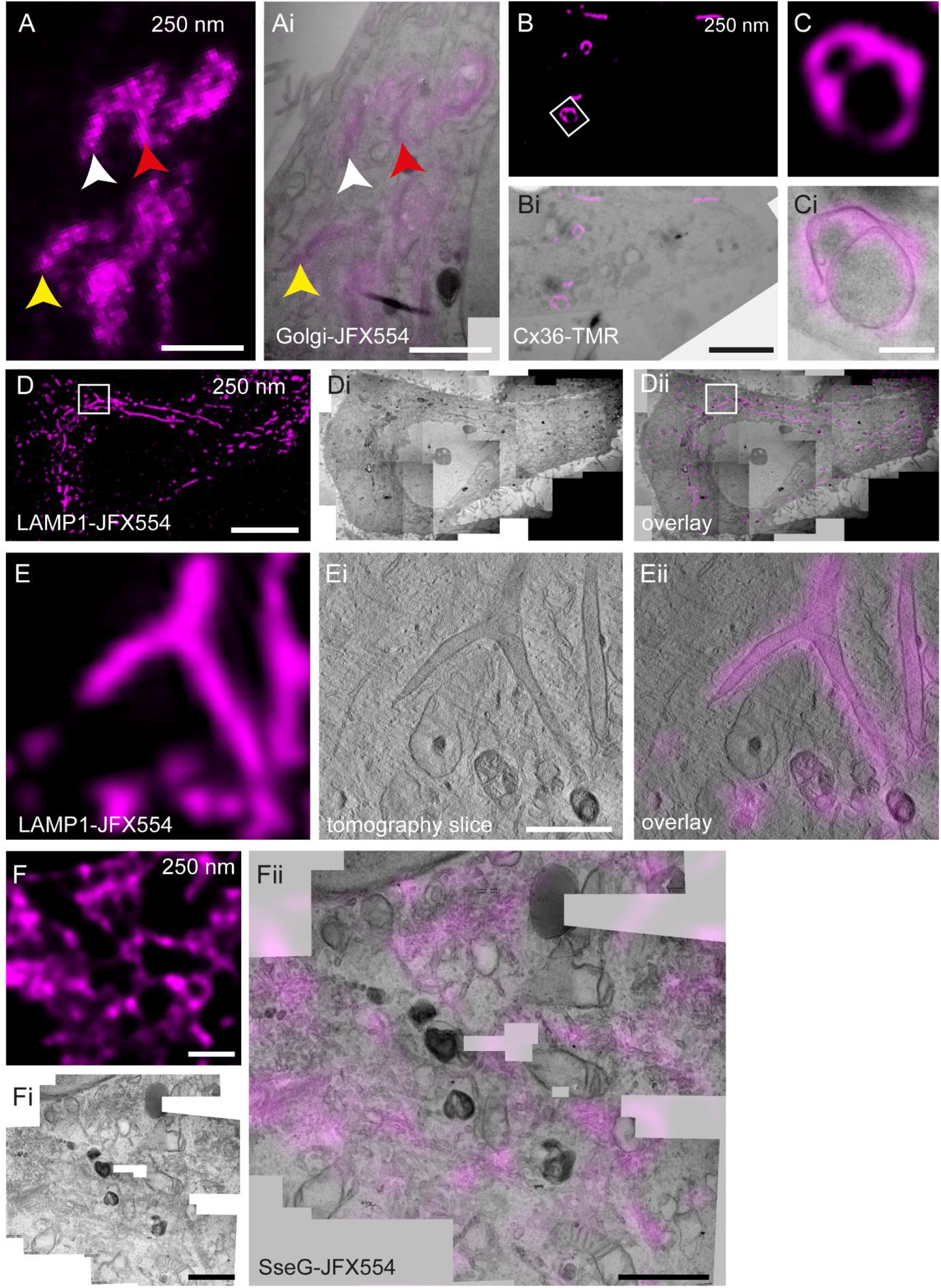
Various cellular structures can be addressed via in-resin CLEM. HeLa (**A**, **D**, **E**, F) or HEK (**B**, **C**) cells were transiently transfected for expression of Golgi-HaloTag (**A**, **Ai**), Connexin36-SNAP (**B**, **Ci**), LAMP1-HaloTag (**D**, **E**), or the *Salmonella* Typhimurium effector protein SseG-HaloTag (**F**), and stained with 100 nm JFX554 or TMR for 30 min. Sections of 250 nm (**A, B, C, D, E**) and 100 nm (**F**) thickness were prepared and imaged by LM (**A, B, C, D, E, F**). Fluorescence signals were retained allowing identification of labelled structures. Next, TEM tomography (**Ai, Ei**) or standard TEM imaging (**Bi, Ci, Fi, Fii**) were performed, and precise correlations of the Golgi apparatus (**Ai**) or ER whorls (**Ci**) were obtained. HeLa cells expressing LAMP1-HaloTag were infected with *S.* Typhimurium stained with JFX554 and fixed 8 h post infection (**D, E,** see Movie 2 for tomogram). Tubular LAMP1-positive structures induced by STM infection are clearly visible. A branching SIF can be observed in LM and TEM tomography (**E, Ei, Eii**). Correlation reveals the tubular architecture of LAMP1-positive membranes (**Eii**). After expression in HeLa cells, SseG is localized at various compartments of the Golgi apparatus (**F, Fi, Fii**). Scale bars: A, Ai: 2 µm; B, Bi: 5 µm; C, Ci: 500 nm; D, Di, Dii: 10 µm; E, Ei, Eii, F, Fi, Fii: 1 µm.

To validate that also further biological questions can be tackled with the in-resin CLEM approach, we set out to recapitulate CLEM results for the endosomal remodeling in mammalian infected by *Salmonella enterica* serovar Typhimurium (STM) (Krieger et al., 2014). For this, HeLa cells expressing LAMP1-HaloTag were infected with STM, at 8 h post infection the cells were stained with JFX554 and prepared for in-resin CLEM. The fluorescence signal of LAMP1-positive *Salmonella*-induced filaments (SIFs) and also other endosomal structures was clearly visible in 250 nm thin EPON sections (**Fig. 5**D). The accurate correlation allowed us to precisely identify a branching filament and to visualize the complicated membrane architecture via TEM tomography (**Fig. 5**E, Ei, Eii). In the same area, also other structures positive for LAMP1 were identified.

To further benchmark the system, we transfected HeLa cells for expression of STM effector protein SseG conjugated with HaloTag, and labelled the fusion protein with JFX554. After sample preparation, the fluorescence signal was still detectable in-resin (**Fig. 5**F) which greatly facilitated correlation of the effector protein with parts of the Golgi apparatus (**Fig. 5**Fii).

These results clearly demonstrate that the rhodamine dyes TMR, JFX549 and JFX554 are suitable markers for targeting of different cellular structures on 250 nm thin EPON sections, while keeping the sample preparation extremely simple. Thus, high precision on-section CLEM can be performed in a fast and convenient way. Routinely incorporating such a workflow into conventional EM labs is perfectly possible and does not require additional equipment or chemicals.

### In-resin Lattice Light Sheet Structured Illumination Microscopy utilizing JFX554

A main problem of CLEM workflows is the large gap in resolution between LM and EM. To overcome this issue, researchers started to integrate super-resolution microscopy (SRM) into CLEM workflows. To evaluate the possibility of using red-fluorescent Janelia Fluor dyes for an in-resin SRM workflow, we deposited 250 nm EPON sections on 50 mesh grids and registered FM modalities using Lattice Light Sheet Microscopy (LLSM) on a set-up also capable of performing Structured Illumination Microscopy (SIM). We selected LLSM because other setups such as Total Internal Reflection Microscopy (TIRF) require very plane surfaces such as glass coverslips to obtain high quality images. However, EPON sections on EM grids are usually not perfectly flat, limiting TIRF imaging. Due to the fact that LLSM does not require completely plane surfaces, we predicted it to be more suited to image the varying surface of the EPON section on the grid. To test this, we labelled HeLa Tom20-HaloTag cells with JFX554, cut sections of 250 nm thickness, deposited them onto a 50 mesh EM grid coated with formvar and inserted it into a custom-made holder for LLSM. With this approach we were able to successfully record in-resin SIM data, which, in comparison to standard LLSM imaging, definitely benefits from the increased resolution allowing us to clearly depict the gap between membrane system and matrix of mitochondria (**Fig. 6**A, B, C, white arrowheads). Finally, retrieval of the same cell in TEM and a tomogram facilitated a precise correlation between both imaging modalities (**Fig. 6**D, E).

**Fig. 6.**
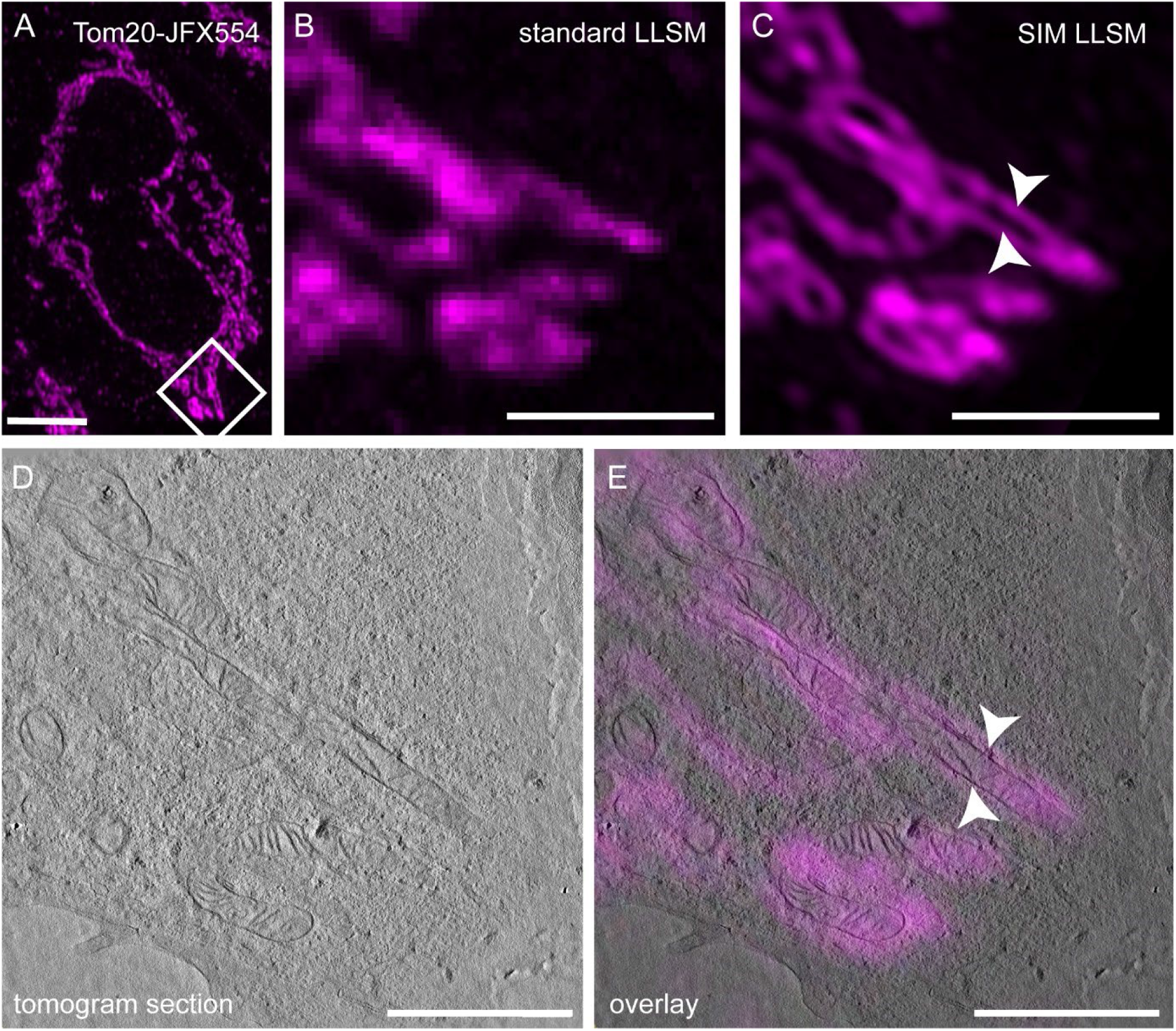
Application of JaneliaFluor HaloTag ligands for in-resin super-resolution microscopy and CLEM. HeLa cells expressing Tom20-HaloTag were labelled with JFX554 as before. 250 nm thin sections were prepared and subjected to lattice light-sheet microscopy (LLSM) in combination with structured illumination microscopy (SIM, **A, B, C**). The comparison of images obtained by LLSM (**B**) or SIM (**C**) modalities indicates the improved spatial resolution of SIM. After performing TEM tomography (**D,** see Movie 3 for tomogram), the super-resolved LM image was precisely correlated with mitochondrial membranes (**E**). Scale bars: A: 5 µm; B, C, D, E: 2 µm.

### In-resin CLEM with HPF and FS

High-pressure freezing (HPF) and freeze substitution (FS) are deployed for preserving ultrastructural features in a closer to native state than conventional EM sample preparation does *(McDonald & Auer, 2006)*. Since the retention of TMR fluorescence after HPF and FS including osmium staining and EPON embedding was previously reporter (Müller et al., 2017), we tested if this is also true for JFX554 by strictly following the protocol from the aforementioned publication. We again utilized HeLa Tom20-HaloTag cells for this to keep results comparable to prior experiments. On 250 nm thin sections of these samples the fluorescence signal of JFX544 was clearly detectable and allowed correlation with the corresponding cell in the TEM (**Fig. 7**A – Aii). Several cellular features such as nucleus, Golgi apparatus and multivesicular bodies were well preserved, and mitochondria within the tomography volume were precisely correlated to JFX554 signals (**Fig. 7**B – Bii). These results indicate that JFX554 performs at least equal to TMR for in-resin CLEM after HPF and FS sample preparation.

**Fig. 7.**
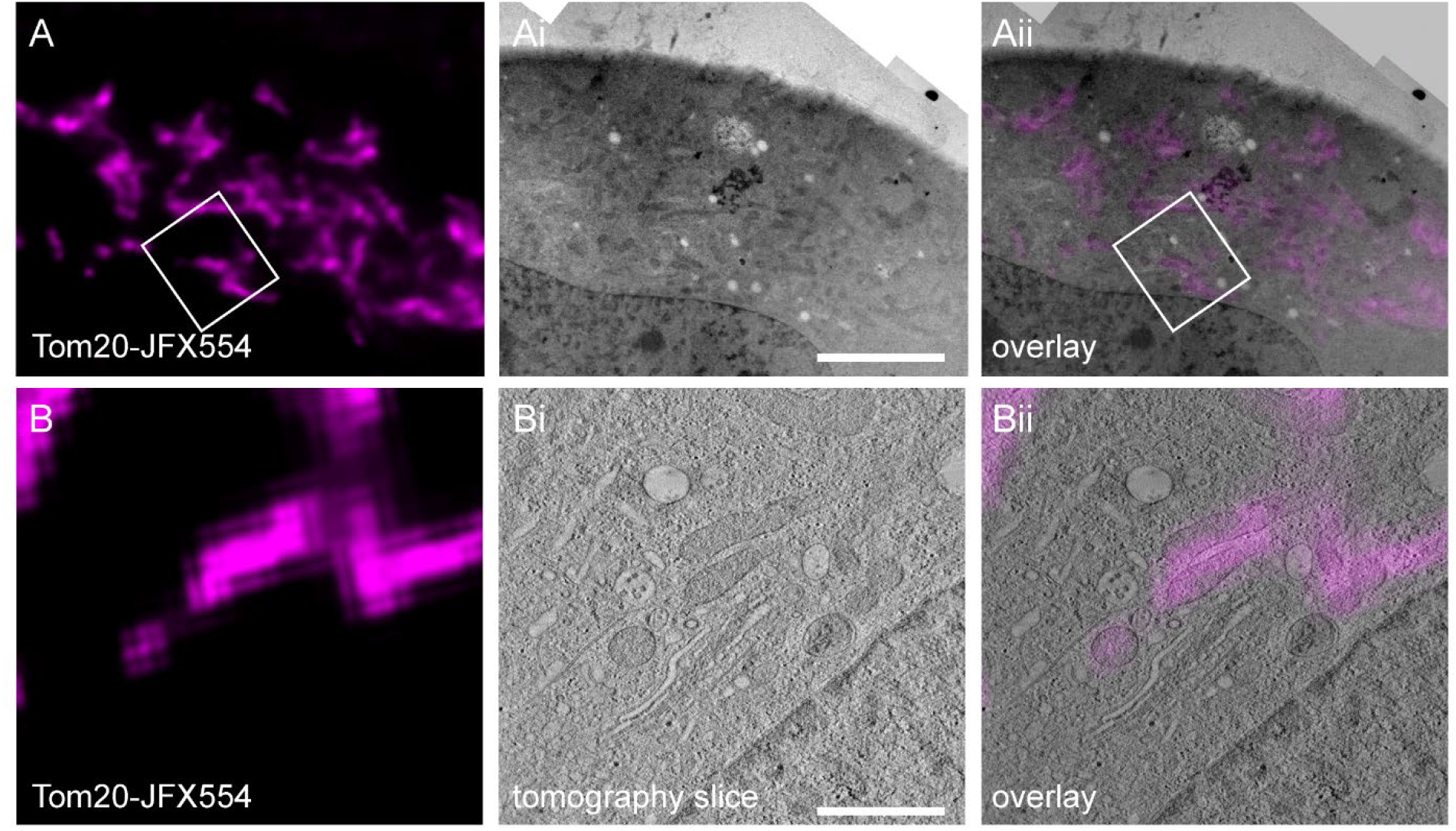
In-resin CLEM after sample preparation by high-pressure freezing and freeze substitution. HeLa cells expressing Tom20-HaloTag were labelled with JFX554 as described before and subjected to HPF and FS. 250 nm sections were observed in LM (**A, B**), TEM (**Ai**) and TEM tomography (**Bi,** Movie 2) modalities. Fluorescence signals were retained after HPF and FS, allowing correlation of labelled mitochondria within the TEM tomogram (**Bii,** see Movie 4 for tomogram). See. Scale bars: A, Ai, Aii: 5 µm; B, Bi, Bii: 1 µm.

## Discussion

Our study examined the performance of selected HTL-conjugated dyes, namely JF479, JF549, JFX549, JFX554, JF585, TMR, JF646, JFX646, JFX650, in a post-embedding in-resin CLEM approach. Initial LM revealed that labeling HeLa Tom20-HaloTag cells with 100 nM of each dye for 30 min before fixation provided sufficient fluorescence signals matching the distribution of mitochondria for most dyes. Only JF479 and JF585 presented an exception, as their fluorescence signal was barely visible in LM. The various fluorophores used here have distinct properties regarding membrane permeability, labeling kinetics, excitation and emission spectra, and may required individual optimization to perform for in-resin CLEM. Here we rather opted to identify fluorophores that can be easily integrated in standard TEM sample preparations and perform with standard microscopy equipment and settings, and in reasonable concentrations. By applying the same labeling conditions for in-resin CLEM studies, we observed in-resin fluorescence only in cells labelled with TMR, JF549, JFX549 and JFX554. All samples were fixed with osmium tetroxide and embedded in EPON. In-resin fluorescence was detectable using the HaloTag or SNAP-tag after labeling structures such as ER, Golgi apparatus, mitochondria, or the *Salmonella* effector protein SseG. Our results demonstrate that a subset of fluorophores retains fluorescence in resin, allowing CLEM applications.

With respect to TMR, the concentration of 100 nM for 30 min used here was considerably lower than in previous work by Los *et al*. (2008) in LM (5 µM TMR for 15-60 min), Perkovic *et al*. (2014) in Lowicryl sections after HPF and FS (50 µM TMR for 30 min), or by Müller *et al*. (2017) in EPON sections chemically fixed and stained with osmium tetroxide (0.6 µM overnight or 6 µM for 1 h using TMR-Star). Using lower fluorophore concentration brings the advantage of lower cell toxicity, reduced preparation costs, and the reduction of washing steps needed to remove excess unbound label. In the quantitative comparison of the in-resin fluorescence of different dyes, we found JFX554 and JFX549 to be the best performing among all tested dyes in terms of signal intensity and S/B ratio. Our findings complement the results of Müller *et al*. (2017), who successfully used SNAP- and CLIP-tag in combination with the respective TMR-ligand to label structures such as insulin or LifeAct in post-embedding CLEM experiments, highlighting the suitability of all three commonly used SLE for CLEM.

Our tests for in-resin fluorescence of the green JF479, the red fluorogenic JF585 and the far-red dyes JF646, JFX646 and JFX650, included higher concentrations of 1 µM for each of these dyes, and even 10 µM for JF479 were tested. Previous studies used these JF dyes for LM of labeled proteins conjugated to HaloTag at concentrations between 100-500 nM (Beatty, 2019; Broadbent et al., 2023; Joshua et al., 2022; Klump et al., 2023; Ricker et al., 2022), sometimes up to 1 µM (Binns et al., 2020; Braselmann et al., 2018). However, in our application the increased concentrations did not improve the results on EPON sections.

Why exactly certain fluorophores perform in our workflow and other fail is unclear. The chemical structure of the fluorophores provides some indications for the different outcomes. All fluorophores retaining fluorescence in-resin possess an oxygen atom within the xanthene structure. In contrast, this atom is replaced by silicon in JF646, JFX646 and JFX650, by nitrogen in JF479, or by carbon in JF585. Thus, this oxygen atom may stabilize the fluorophore during EM sample preparation. To test this hypothesis, the dye JF525 (Grimm, Muthusamy, et al., 2017) could be examined, as it features the same structure as JF479 and JF585, but possesses an oxygen atom in the xanthene instead of nitrogen or carbon atoms, respectively. Alternatively, the difference between the fluorophores could arise solely from the silicon and the 3,3-difluoroazetidines as these two features apply only to the far-red dyes and to JF479 and JF585, which lost in-resin fluorescence. In order to verify this hypothesis at least with respect to the 3,3-difluoroazetidines, the antecedents of JF479 and JF585, namely JF502 and JF608 (Grimm, Muthusamy, et al., 2017; Grimm et al., 2020), could be tested as they comprise nitrogen and carbon atoms, respectively, but not the 3,3-difluoroazetidines.

To identify and optimize fluorophores for in-resin CLEM, future work should systematically investigate the effect of various modifications of the JF dyes chemical structures on their resistance to conventional EM sample preparation. In this regard dyes may be engineered which are especially resistant and useful for in-resin CLEM similar to optimization of fluorescent proteins (FP) for EM such as mEosEM (Fu et al., 2020). Moreover, optimized variants of SLE with increased fluorescence intensity of bound dyes should be tested for in-resin fluorescence retainment (Frei et al., 2022).

A recent approach to simplify post-embedding CLEM deploys specifically engineered FP which can, to some degree, tolerate the quenching properties of a conventional sample preparation protocol (Fu et al., 2020; Paez-Segala et al., 2015; Peng et al., 2022; Tanida et al., 2020). However, many of these FP apparently retain only comparably weak fluorescence signals and low S/B ratio after EPON embedding. For sufficient preservation of FP fluorescence, reduced osmium tetroxide concentrations and/or incubation times such as 10 min were applied. This, however might severely reduce ultrastructural details in TEM modalities. Furthermore, in these publications, in-resin fluorescence was always registered on sections being on glass coverslips. This format allows SEM imaging, but is not compatible with 2D or 3D TEM imaging, or requires the removal of the glass from the resin sections and support film via hazardous hydrofluoric acid etching (Fu et al., 2020; Paez-Segala et al., 2015; Peng et al., 2022; Sanada et al., 2022; Tanida et al., 2020).

The importance of testing the performance of various dyes in on-section CLEM workflows with conventional sample preparation and EPON embedding is supported by recent publications in this field (Müller et al., 2017; Sanada et al., 2022; Tanida et al., 2023). So far, only Müller *et al*. worked with cell-permeable dyes in such setup, and reported successful application of TMR and 505-Star coupled to ligands of HaloTag, SNAP-tag or CLIP-tag. However, the protocols used in Müller et al. (2017) calls for strongly reduced osmium concentrations of 0.1%, and did not deploy potassium hexacyanoferrate as a further contrast enhancer, as done in our approach. Since their study was not focused on systematically or quantitatively testing the performance of a larger set of dyes, we here aimed to tackle this point. Interestingly, the group showed that 505-Star within insulin granules survived sample preparation. We speculate that either large amounts of dye accumulated in these insulin granules and were not completely quenched during the sample preparation, or that 505-Star represents a promising alternative to the green dyes tested in our work. This could conceivably be the case, because the structure of 505-Star contains an oxygen atom within the xanthene, comparable to JF525 or JFX554, and following our earlier hypothesis, possibly making it more resistant to EM sample preparation. Sanada *et al*. (2022) and Tanida *et al*. (2023) investigated various cell-impermeable dyes for their performance after conventional sample preparation and EPON embedding and found that red dyes such as DyLight549, iFluor546 or HiLyte-555 performed best, while detection of Alexa Fluor 647 was also still possible in resin. Our results with cell-permeable dyes complement these findings, as we also identified red dyes as best-performing, while Alexa Fluor 647 also retained fluorescence. A great downside of the protocol employed in their work (Sanada et al., 2022; Tanida et al., 2023) is the need for cell permeabilization to allow dyes to access intracellular targets. Permeabilization strongly affects ultrastructural quality of EM samples (Humbel et al., 1998).

A comparable, well-established method to preserve the fluorescence within biological samples consists of embedding high-pressure frozen (HPF) samples in Lowicryl resins after freeze substitution (FS) with low concentrations of uranyl acetate (Ader et al., 2019; Buerger et al., 2021; Kukulski et al., 2011; Perkovic et al., 2014). For this, no special dyes or SLE are needed, but standard fluorescent proteins like GFP or mCherry can be used. For some JF dyes, we also demonstrated the applicability in HPF/FS workflows with EPON embedding, and we anticipate that dyes will also perform at least as equally well in hydrophilic resins, or at better labeling conditions. However, such workflows are challenging, require advanced and expensive equipment and osmium tetroxide cannot be used to enhance contrast for EM. Furthermore, the best ultrastructural preservation is most often obtained in EPON resin (Kann & Fouquet, 1989). Therefore, the workflow described here provides an easy and cheap alternative for in-resin CLEM which can be incorporated by every EM facility without need for additional equipment or fundamental changes in sample preparation protocols to ensure fluorescence preservation. In addition, the here tested JF dyes allow in-resin SRM improving the correlation of LM and EM modalities (Sims & Hardin, 2007).

In conclusion, our analyses provide a first systematic basis to further characterize und understand the behavior of fluorophores after conventional sample preparation for TEM. The red dyes JFX554 and JFX549 represent the most promising candidates for use in conventional in-resin CLEM approaches. We hypothesize that specifically the oxygen atom within the xanthene structure of the various dyes plays an important role in withstanding the harsh conditions of the protocol, and in retaining fluorescent properties to sufficient degree. Overall, the workflow described here has the advantages of being fast, easy to use, only requiring low concentrations of dyes, and being easily integrated into standard workflows of imaging facilities.

## Materials and Methods

### Cell lines and cell culture conditions

The epithelial cell line HeLa (American Type Culture Collection, ATCC no. CCL-2) and derivatives (Table 2) were cultured in Dulbecco’s modified Eagle medium (DMEM) containing 4.5 g/l glucose, 4 mM stable glutamine and sodium pyruvate (Biochrom), and 10% inactivated fetal calf serum (iFCS) (Sigma-Aldrich) at 37 °C in an atmosphere containing 5% CO_2_ and 90% humidity. HEK-293FT (humane embryonic kidney, Invitrogen no. R700-07) cells were cultured the same way. For CLEM, cells were seeded into 8-well µ-slides with polymer bottom (ibidi) with (Art. 80826-G500) or without (Art. 80806) an engraved coordinate system. Cell culture was performed to achieve about 50-60% confluency on the day of the experiment.

**Table 1.**
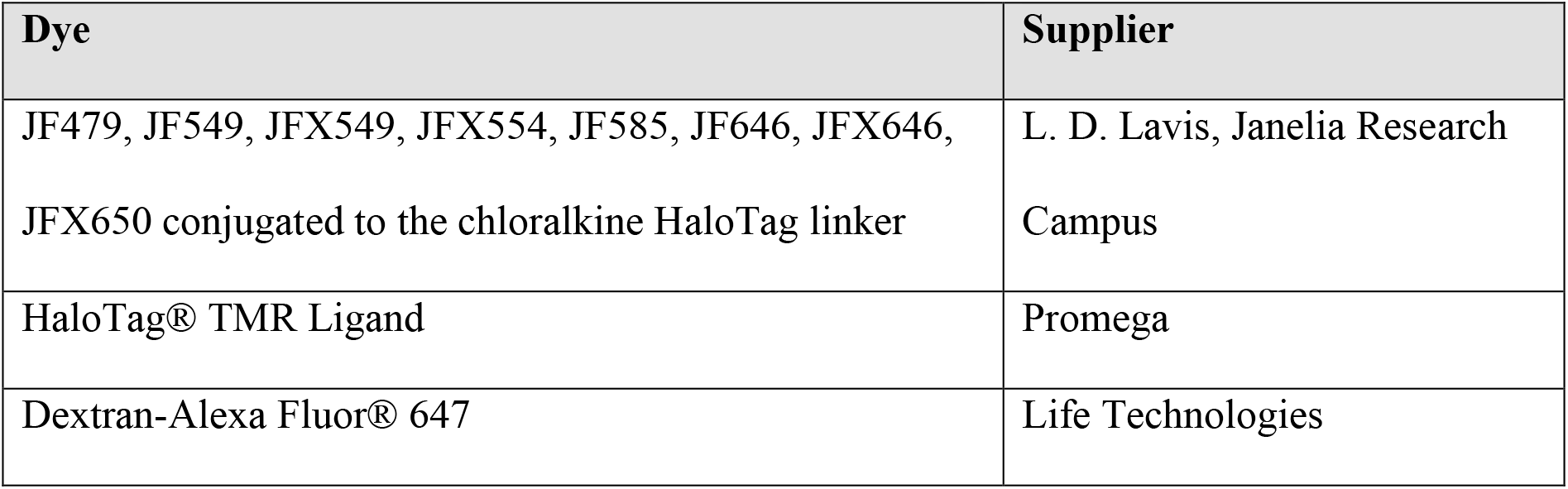
Dyes used for staining.

**Table 2.** Cell lines used in this study.

| **Designation** | **relevant characteristics** | **source/reference** |
| --- | --- | --- |
| HeLa Tom20-HaloTag | HeLa WT cells, permanent transfected with Tom20-HaloTag | AG Piehler, Biophysics UOS |
| HeLa LAMP1-HaloTag | HeLa WT cells, permanent transfected with LAMP1-HaloTag | AG Hensel, Microbiology, UOS |
| HeLa LAMP1-GFP | HeLa WT cells, permanent transfected with LAMP1-GFP | AG Hensel, Microbiology, UOS |

### Transfection of cells

HeLa or HEK cells were cultured for at least one day and transfected using FUGENE HD reagent (Promega) according to manufacturer’s instruction. Plasmids used for transfection are listed in Table 3. Briefly, 0.5–2 µg of plasmid DNA were solved in 25–100 µl DMEM without iFCS and mixed with 1–4 µl FUGENE reagent (ratio of 1:2 for DNA to FUGENE). After 10 min incubation at room temperature (RT) the transfection mix was added to the cells in DMEM with 10% iFCS for at least 18 h. Before infection or staining, cells were provided with fresh medium without transfection mix.

**Table 3.** Plasmids used in this study.

| **Designation** | **Purpose** | **genotype** | **source/reference** |
| --- | --- | --- | --- |
| pFPV25.1 | bacterial expression | $P_{rpsM}::eGFP$ mut3 | AG Hensel,<br>Microbiology, UOS |
| p4564 | mammalian expression | *sseG*::HaloTag | AG Hensel,<br>Microbiology, UOS |
| p6032 | mammalian expression | Golgi::HaloTag | AG Hensel,<br>Microbiology, UOS |
| pCx36-SNAP | mammalian expression | Cx36::SNAP-tag | (Tetenborg et al., 2023) |

### Infection of cells

For infection of HeLa cells, *Salmonella enterica* serovar Typhimurium NCTC12023 strains were grown in LB broth with appropriate antibiotics overnight (O/N), diluted 1:31 in fresh LB with antibiotics and subcultured for 3.5 h at 37 °C. The infection of HeLa cells was performed at various multiplicities of infection (MOI) for 25 min by directly adding a suitable amount of *Salmonella* subculture to the cells. Subsequently, cells were washed thrice with PBS and incubated for 1 h with DMEM containing 100 µg/ml gentamicin (Applichem) to kill extracellular bacteria. Finally, the medium was replaced by DMEM containing 10 µg/ml gentamicin for the rest of the experiment.

### Staining of cells with fluorescently labelled ligands

Dyes coupled to their respective ligands as listed in Table 1 were solved in DMSO, diluted in PBS or cell culture medium, and directly added to cell culture medium yielding the appropriate final concentrations. After 30 min of incubation time at 37 °C, 5% CO_2_, 90% humidity, if not mentioned otherwise, stained cells were washed with PBS 3-5 x for 1 min to remove unbound dye, and subsequently fixed.

### Fixation of cells for pre-embedding light microscopy

For pre-embedding light microscopy, 37 °C pre-warmed fixative containing 3% PFA and 0.1-0.2% GA in 0.1 M sodium cacodylate buffer (pH 7.2) was added to cells and incubated for 30 min at RT. Subsequently, samples were washed 3-5x for 1 min with 0.1 M sodium cacodylate buffer and either directly imaged or stored in the dark at 4 °C.

### EM sample preparation

During the entire procedure, samples were kept in the dark as far as possible. First, samples were fixed with either pre-warmed (living cells) or 4 °C cold (fixed cells for LM) 2.5% GA in 0.1 M sodium cacodylate buffer (pH 7.2) for 1 h. Living cells were kept for 15 min at 37 °C in fixative and were then transferred to RT for further incubation. Then, samples were placed on ice for another 30 min before washing with 0.1 M sodium cacodylate buffer (pH 7.2) thrice for 1 min. Samples were post-fixed for 30 min with 1% OsO_4_ + 1.5% K_4_[Fe(CN)_6_] in 0.1 M sodium cacodylate buffer (pH 7.2). Subsequently, samples were washed 5x for 1 min with 0.1 M sodium cacodylate buffer (pH 7.2) and dehydrated. For dehydration, samples were successively incubated in 30%, 50% 70%, 80%, 90%, 100% ethanol and 100% anhydrous ethanol for 7 min each on ice and allowed to reach RT during the last step. After two incubation steps in 100% anhydrous acetone for 7 min each, samples were infiltrated on a shaker (50-60 rpm) with 25%, 50%, and 75% EPON (Sigma-Aldrich) diluted in anhydrous acetone for 1 h each with closed lids. Lids were removed from 8-well µ-slides and the samples were kept on the shaker in 100% EPON O/N. EPON was exchanged the next morning and again after 6 h. Thereafter polymerization was conducted at 60 °C for 48 h. The polymerized EPON blocks were sawed out from the 8-well µ-slides together with the polymer bottom which was removed by placing the bottom of the EPON blocks in toluol (Roth) and frequently wiping it with a paper towel. Each EPON block was sawed into 4 smaller blocks, trimmed to a trapezoid and sectioned into 100 nm or 250 nm sections using an ultramicrotome (Leica ULTRACUT EM UC7RT or UC7cryo). For quantitative comparison experiments the sections were placed on a glass coverslip previously washed in 100% ethanol. For correlative experiments the sections were transferred to carbon- or formvar-coated EM grids.

### Pre-embedding light microscopy

Light microscopy prior to EM sample preparation was either conducted at a Leica CLSM SP5 equipped with HC PL FL 10x (NA 0.3), HCX PL APO CS 40x (NA 0.75 – 1.25, oil immersion) and HCX PL APO CS 100x (NA 0.7 – 1.4, oil immersion) objectives or at a Zeiss Cell Observer SD equipped with Alpha Plan-Apochromat 63x (NA 1.46, oil immersion) and Plan-Apochromat 10x (NA 0.45, air) objectives. Samples were screened for ROIs and subsequently high magnification Z-stacks (100x or 63x objectives) and low magnification (10x objectives) overview images revealing the 8-well µ-slide coordinate system for relocation were acquired.

### Light microscopy of EPON sections (in-resin LM)

For in-resin LM, a 10 µl drop of PBS (pH 8.4) was added respectively onto two 25 mm glass coverslips. The EM grid carrying the resin sections was placed on top of one of the PBS drops and subsequently sandwiched with the remaining coverslip. The assembly was transferred into a custom-made holder and imaged with the sections facing the objective lens using an Olympus FV-3000 with settings listed in Table 5. The microscope was either operated as a CLSM or as a widefield system as indicated in figure legends and was equipped with a sCMOS camera (ORCA-Flash 4.0, Hamamatsu, Japan). Using appropriate filter/detector settings for the specific dyes, Z-stacks (step-size 200-400 nm) were acquired with a 60x oil immersion lens (PLAPON-SC NA 1.4). Subsequently, overview images of the grid utilizing the 10x air objective (UPL SAPO NA 0.4) facilitating correlation later on in the TEM were recorded. Frequently used exposure times for visualizing the different dyes ranged between 500 ms – 1000 ms. After imaging, the grid was removed from the coverslip sandwich, washed thrice in distilled water, dried by touching a filter paper and stored in the dark.

**Table 4.**
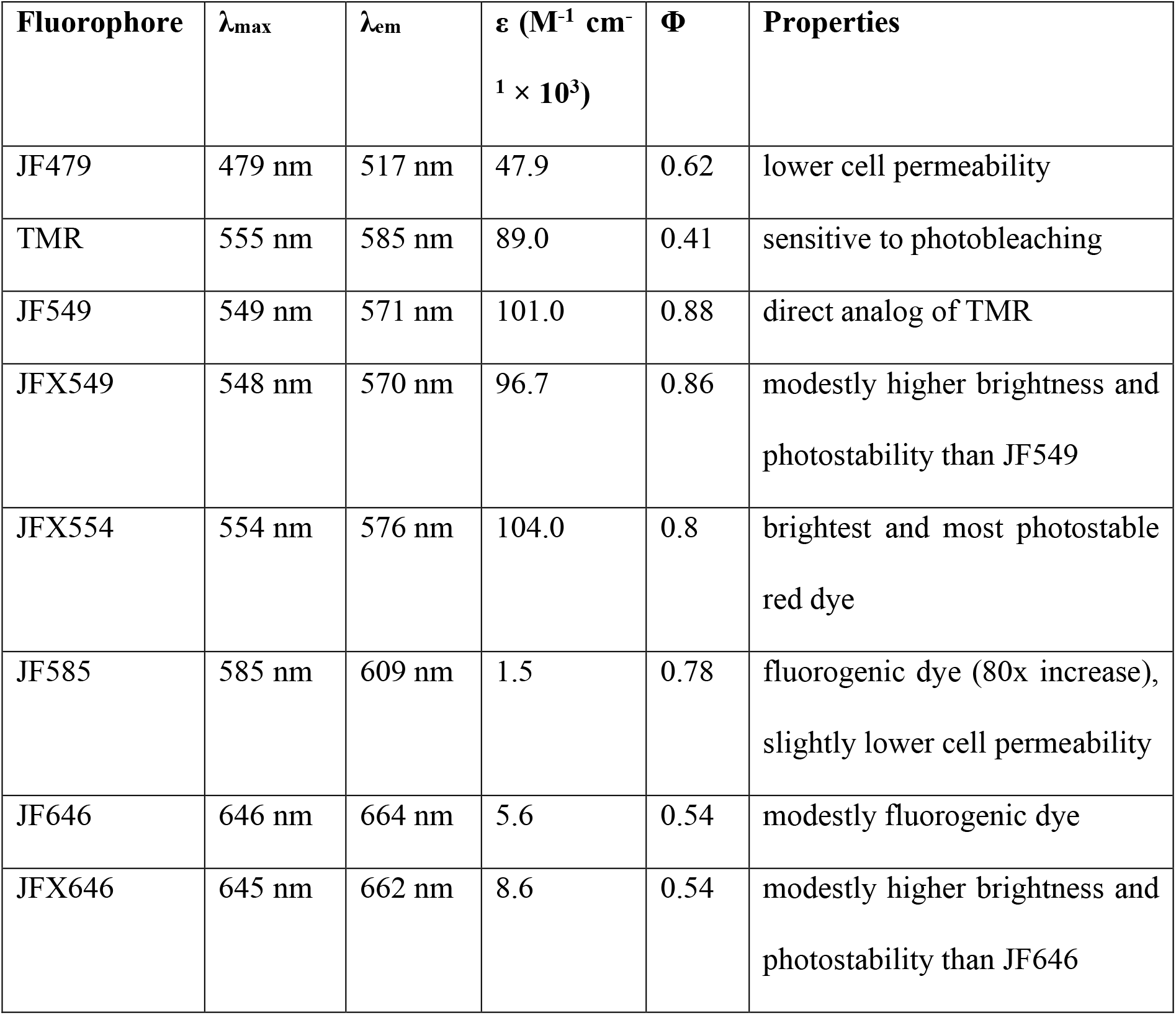
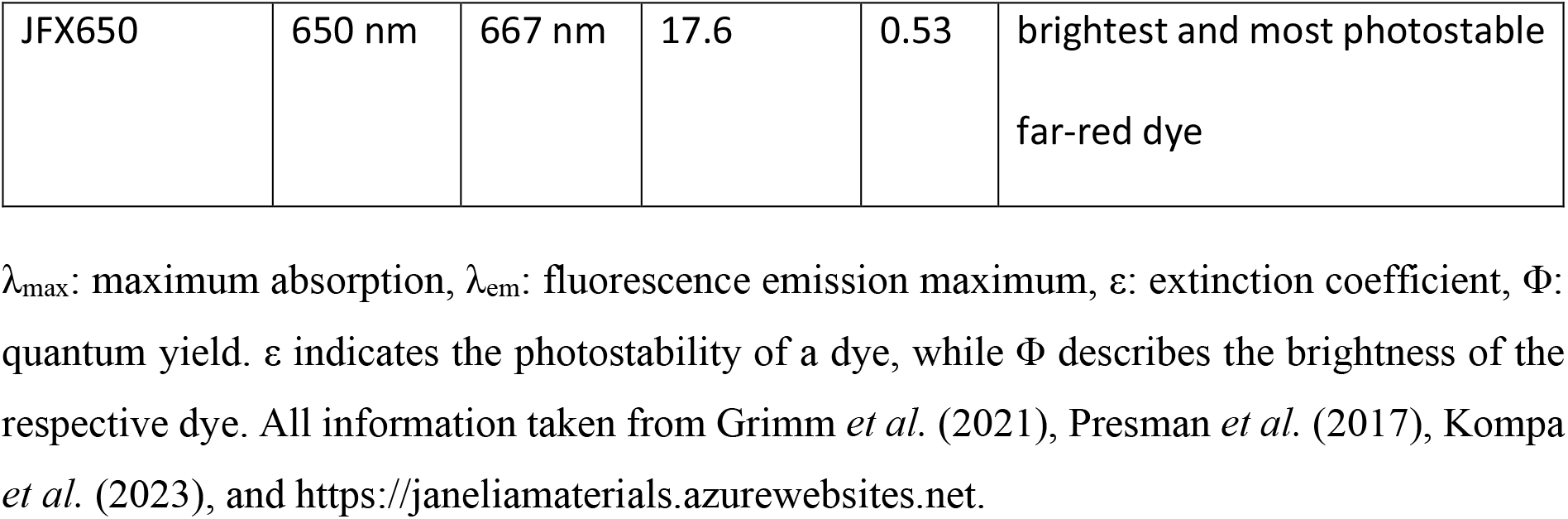
Physical properties of the fluorophores analyzed in this study.

**Table 5.** Settings for quantitative image acquisition at Olympus LSM FV3000 NLO.

| Dye | Channel | Excitation<br>wavelength | LED | Exposure<br>time | Cond<br>ensor | Filter |
| --- | --- | --- | --- | --- | --- | --- |
| JF479 | GFP | 488 nm | 100% | 1 s | No | 409/493/573/6<br>52 (LED) |
| JF549, JFX549,<br>JFX554, JF585,<br>TMR | TMR | 560 nm | 30% | 500 m<br>s | No | 409/493/573/6<br>52 (LED) |
| JF646, JFX646,<br>JFX650, Dextran-Alexa Fluor® 647 | Cy5 | 640 nm | 100% | 1 s | No | 409/493/573/6<br>52 (LED) |

### Structured Illumination Microscopy (SIM) using lattice-light sheet microscopy

Lattice light-sheet microscopy was performed on a home-built clone of the original design by the Eric Betzig group (Chen et al., 2014). The EM grid carrying the section was mounted in a sample holder, which was attached on top of a sample piezo. This ensures that the sample is inserted at the correct position inside the sample bath, containing PBS (pH 7.4) at 25 °C. An image stack was acquired in sample scan mode by scanning the sample through a fixed light sheet with a step size of 400 nm which is equivalent to a ∼ 216 nm slicing with respect to the Z axis considering the sample scan angle of 32.8°. We used a dithered square lattice pattern generated by multiple Bessel beams using an inner and outer numerical aperture of the excitation objective of 0.48 and 0.55, respectively. JFX554 was excited using a 561 nm laser (2RU-VFL-P-2000-561-B1R; MPB Communications Inc.). Fluorescence was detected by a sCMOS camera (ORCA Flash 4.0, Hamamatsu, Japan) using an exposure time of 50 ms for each channel. The raw data were further processed by using an open-source LLSM post-processing utility called LLSpy (https://github.com/tlambert03/LLSpy) for de-skewing and deconvolution. Deconvolution was performed by using experimental point spread functions and is based on the Richardson–Lucy algorithm using 10 iterations.

### Transmission electron microscopy (TEM)

Resin sections on EM grids contrasted with 3% uranyl acetate for 30 min and 2% lead citrate for 20 min in the LEICA EM AC20 were analyzed in a JEOL TEM 200 keV JEM2100-Plus equipped with a 20 megapixel EMSIS Xarosa CMOS camera (EMSIS, Münster, Germany), or a Zeiss Leo Omega AB equipped with a CCD camera (Tröndle). To facilitate correlation, first low magnification overview images were recorded and compared to LM overview images. After identifying the ROI, high magnification images of corresponding cells were recorded.

### TEM tomography acquisition

Prior to tomogram acquisition and contrasting, sections on grids were labelled with 10 or 15 nm protein-A gold (PAG) fiducials on both sides. For this, grids were incubated for 3 min on a 1:50 diluted drop of PAG with distilled water and subsequently washed thrice for 1 min with distilled water. Tilt series were acquired from +60° to -60° with 1° increments using the TEMography software (JEOL, Freising, Germany) on a JEM 2100-Plus system operating at 200 keV and equipped with a 20 megapixel CMOS camera. Tomograms were reconstructed using the back projection algorithm in IMOD (Kremer et al., 1996).

### Quantification of fluorescence in EPON sections and statistical analysis

Using the software FIJI, the LM modalities of various dyes were analyzed for quantitative comparison. The custom-made algorithm enhanced the contrast of the image, applied a Gaussian blur and subtracted the background before generating a mask by thresholding. Values measured of the area outside the selection were ascribed to the background. After manually deselecting areas with bright fluorescence, which obviously did not correspond to mitochondria but were rather induced by dust particles or folds in the section, the values of the remaining selection were measured. Lastly, cells were counted manually and the selections were saved as a separate image allowing subsequent clarification if necessary . If the selection clearly traced the mitochondria, the acquired data was further taken into account for analysis. If the algorithm randomly selected various patches over the entire image, the image was considered as not showing any fluorescence. Selections not selecting fluorescence signals within the cells or selecting many background speckles were excluded from further analysis.

## Acknowledgements

This work was supported by the DFG though grants SFB 944, project Z and SFB 1557, project Z2, and the iBiOs platform in priority programme SPP 2225 ‘Exit Strategies’, and instrumentation grants INST 190/180-1 FUGB and INST 190/181-1 FUGB. We thank Luke D. Lavis, Janelia Research Campus, for the generous gift of HaloTag Janelia Fluor ligands.

## Supplemental Movies Captions

Movie 1: Tomogram corresponding to **Fig. 4**B. For download, use https://myshare.uni-osnabrueck.de/f/db35d73b0ebf4ae3b8e0/

Movie 2: Tomogram corresponding to **Fig. 5**E. For download, use https://myshare.uni-osnabrueck.de/f/3cfa267d21dc4144a7b1/

Movie 3: Tomogram corresponding to **Fig. 6**D. For download, use https://myshare.uni-osnabrueck.de/f/e911ac20023649c9be78/

Movie 4: Tomogram corresponding to **Fig. 7**B. For download, use https://myshare.uni-osnabrueck.de/f/8ef1391cd9e64614bf93/

Movie 5: LM Z stack of EPON-embedded cells. The movie corresponds to data shown in **Fig. S 5**A. For download, use https://myshare.uni-osnabrueck.de/f/c521611d9a8348b391f9/

## Supplemental Code

Source code of the algorithm used for quantitative comparison experiments

The source code below can be saved as a .ijm-file, implemented into Fiji under the menu bar Plugins>Macros>Install…, and used for measuring the fluorescence signal of mitochondria.

macro “Auswertung in-resin [q]” {

//get relevant image information

name=getTitle();

directory = File.directory;

//generate mask in a seperate window

run(“Duplicate…”, ” ”);

run(“Enhance Contrast…”, “saturated=0 normalize”);

run(“Gaussian Blur…”, “sigma=3”);

run(“Subtract Background…”, “rolling=50”); setAutoThreshold(“IsoData dark”);

setOption(“BlackBackground”, false);

run(“Convert to Mask”);

run(“Create Selection”);

//transfer selection to original window

selectWindow(name);

//setTool(“rectangle”);

run(“Select None”);

run(“Restore Selection”);

//measure background values

run(“Make Inverse”);

run(“Set Measurements…”, “area mean standard min integrated median display redirect=None decimal=3”);

run(“Measure”);

backarea=getResult(“Area”, nResults-1);

backmean=getResult(“Mean”, nResults-1);

backstddev=getResult(“StdDev”, nResults-1);

backmin=getResult(“Min”, nResults-1);

backmax=getResult(“Max”, nResults-1);

backintden=getResult(“IntDen”, nResults-1);

backmedian=getResult(“Median”, nResults-1);

backrawintden=getResult(“RawIntDen”, nResults-1);

run(“Make Inverse”);

//manually deselect areas of unwanted autofluorescence (created by dirt particles or by coverslip background)

setTool(“brush”);

waitForUser(“Brush”, “Please remove dirt… \nTo adjust size of brush: double click on the brush tool in the menu”);

roiManager(“add”);

//measure fluorescence signal in selected area

run(“Set Measurements…”, “area mean standard min integrated median display redirect=None decimal=3”);

run(“Measure”);

setResult(“background area”,nResults-1,backarea);

setResult(“background mean”,nResults-1,backmean);

setResult(“background StdDev”,nResults-1,backstddev);

setResult(“background Min”,nResults-1,backmin);

setResult(“background Max”,nResults-1,backmax);

setResult(“background IntDen”,nResults-1,backintden);

setResult(“background Median”,nResults-1,backmedian);

setResult(“background RawIntDen”,nResults-1,backrawintden);

setResult(“Label”,nResults-1,name);

IJ.deleteRows(nResults-2, nResults-2);

//manually count cells

setTool(“multipoint”);

waitForUser(“count cells”, “Please mark all cells”);

getSelectionCoordinates(xCoordinates, yCoordinates);

count=lengthOf(xCoordinates);

setResult(“number of cells”, nResults-1, count);

//save image with selection as .jpeg

roiManager(“add”);

roiManager(“Select”, 0);

roiManager(“Set Color”, “yellow”);

roiManager(“Select”, 1);

roiManager(“Set Color”, “red”);

roiManager(“deselect”);

roiManager(“show all”);

resetMinAndMax();

run(“Capture Image”);

print(directory);

print(name);

saveAs(“Jpeg”, directory+name+”_selection.jpeg”);

close();

close(name);

close(substring(name,0,indexOf(name,”.tif”))+”-1.tif”);

roiManager(“deselect”);

roiManager(“delete”);

}

**Fig. S1.**
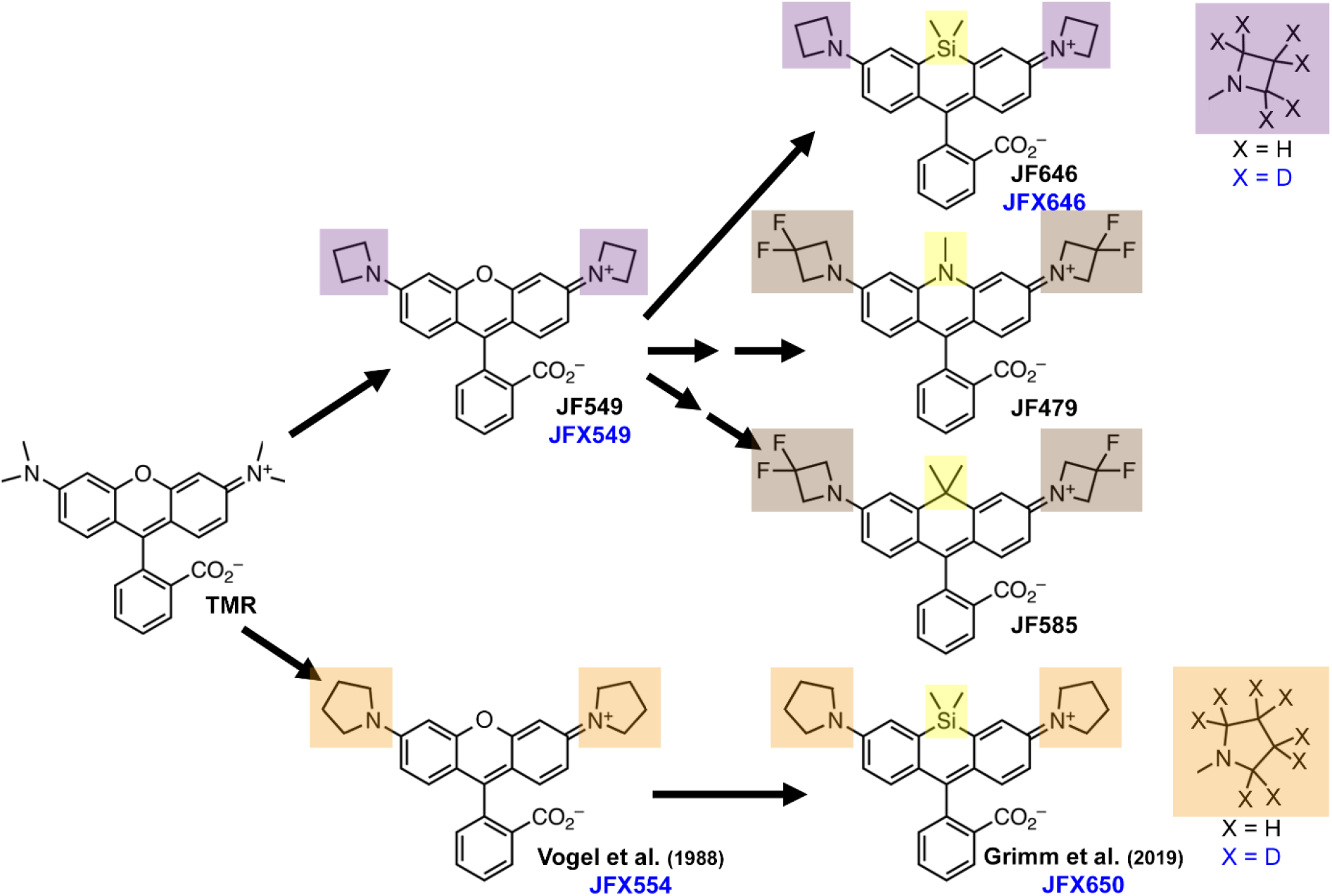
Origin of rhodamine-derived Janelia Fluor dyes. Substitution of the N-alkyl groups of tetramethylrhodamine (TMR) by azetidines (purple), pyrrolidine (orange) or a 3,3-difluoroazetidines (brown) together with replacing the oxygen atom of the xanthene by Si, N or C atoms (yellow) resulted in various dyes. Additional substitution of the hydrogen (H) by deuterium (D) in the azetidines and pyrrolidines improved the fluorescent characteristics of the dyes. The deuterated dyes are marked in blue, while the parent structures are marked in black. Structures shown were adapted from Janelia Materials (2023).

**Fig. S2.**
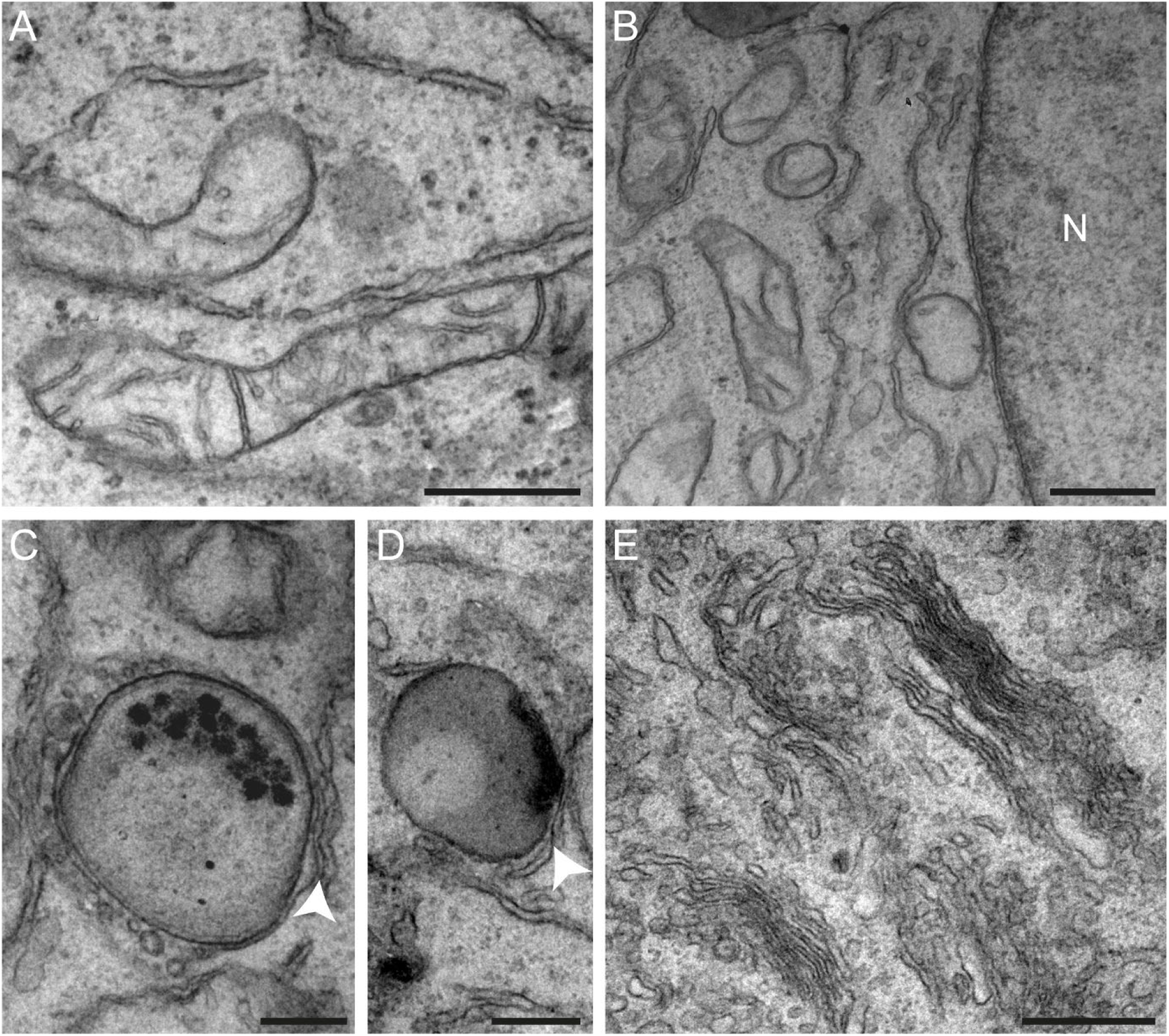
Ultrastructural preservation after OsO_4_ treatment for 30 min. HeLa cells were embedded as described in Material and Methods. Ultrathin sections were prepared to ensure good ultrastructural preservation even after the reduced duration of OsO_4_ staining. Several cellular structures including mitochondria and ER (**A**), Nucleus (N) with double membrane (**B**), different stages of lysosomes and membrane contact sites (MCS) with ER (**C, D**), as well as parts of the Golgi apparatus (**E**) are well preserved. Arrowheads indicate potential MCS. Scale bars: A, B, E: 500 nm; C, D: 250 nm.

**Fig. S3.**
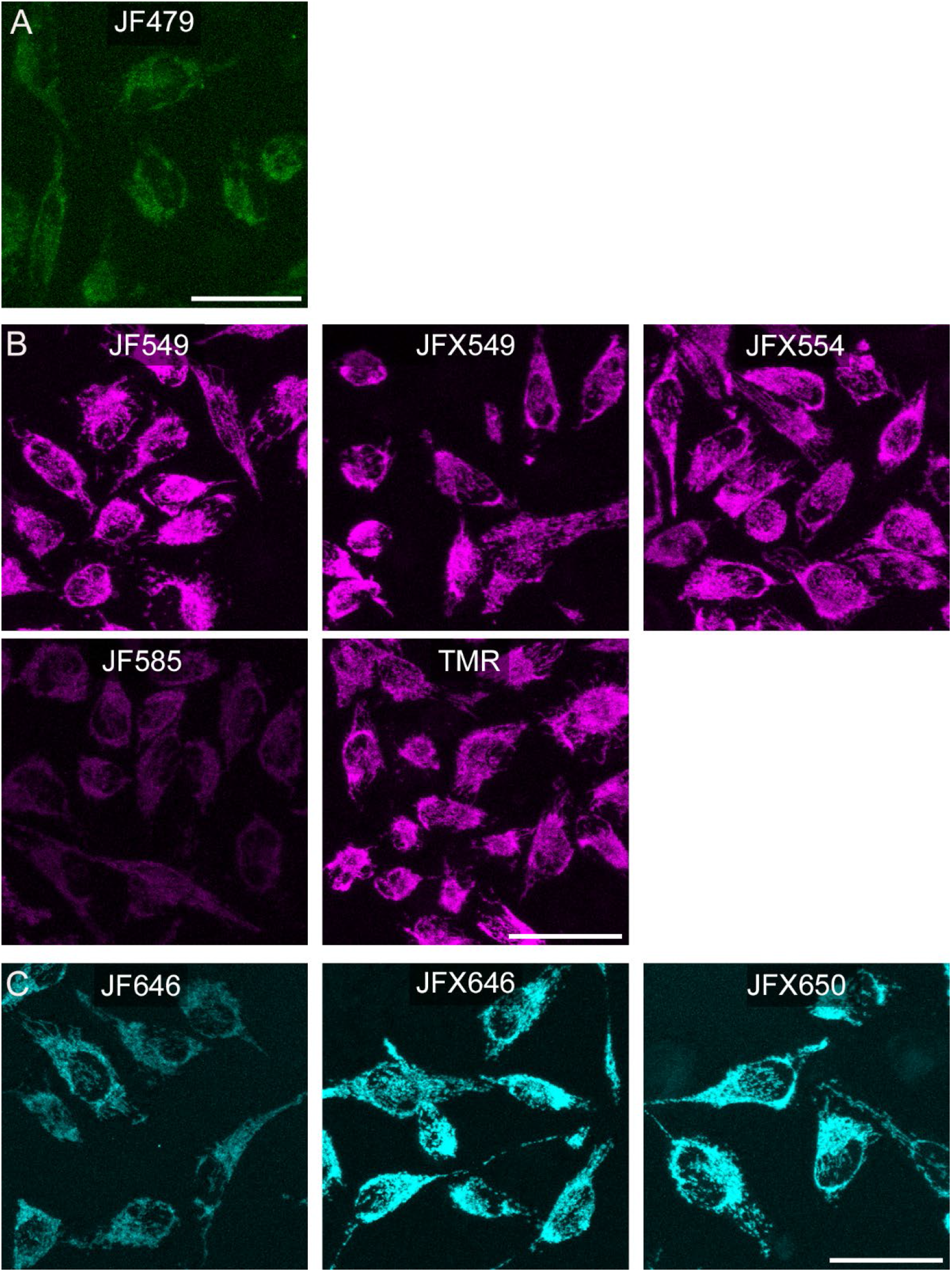
Specific labelling and sufficient fluorescence signal intensities were observed for all dyes tested. HeLa Tom20-HaloTag cells were stained with 100 nM of the indicated **A**) green, **B**) red, or **C**) far-red fluorescent dyes for 30 min. Fixation was performed with 3% PFA for 15 min. Image stacks of the same size were acquired using confocal laser scanning microscopy (Leica SP5) with the same settings in the respective channel and were later processed the same way using maximum intensity projections. Scale bars: 50 µm.

**Fig. S4.**
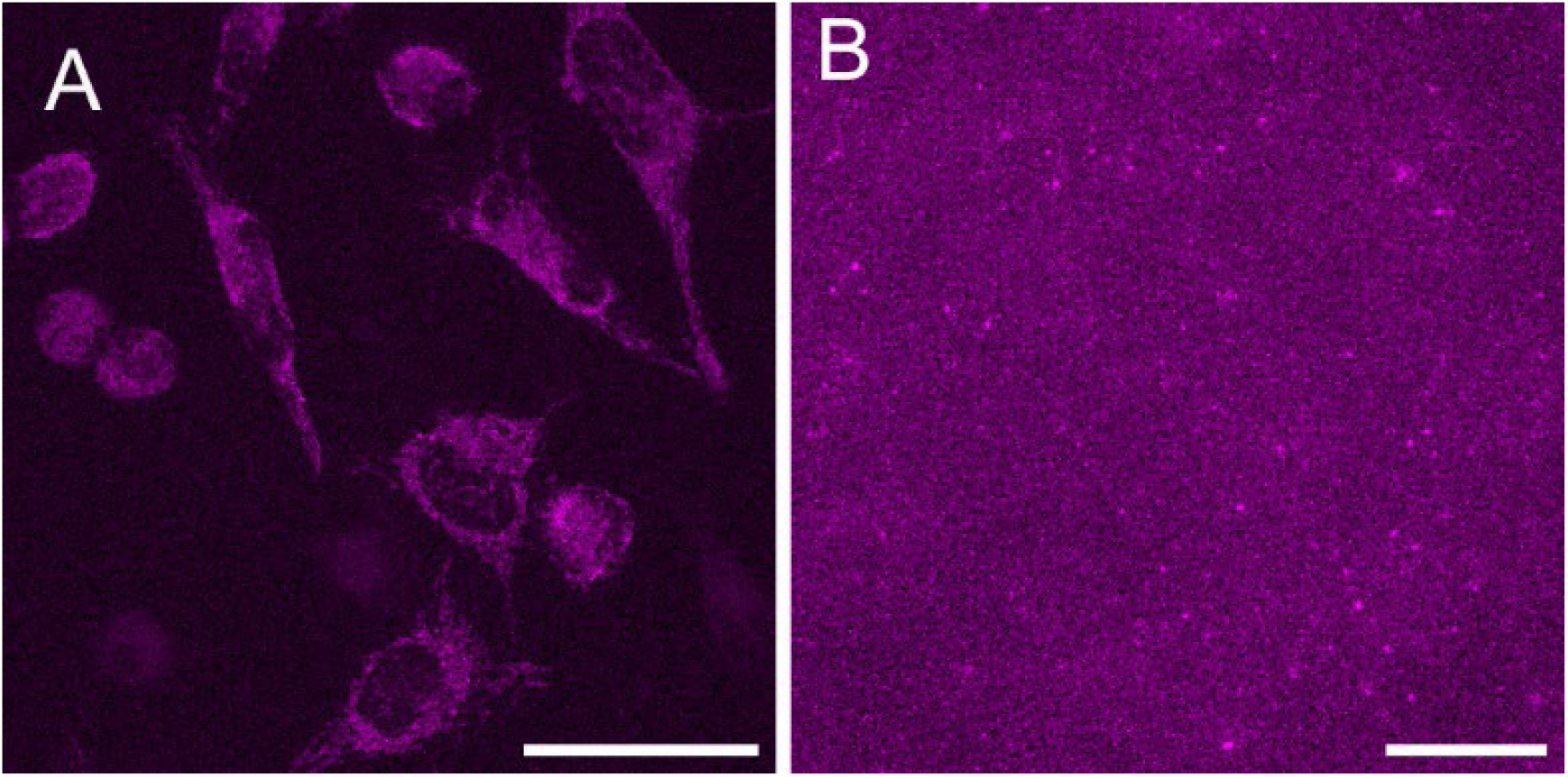
In-resin fluorescence is not retained for JF585 at a higher concentration of 1 µM. **A**) Fluorescence image after 30 min of time lapse microscopy of living HeLa Tom20-HaloTag cells stained with 1 µM JF585. **B**) A 250 nm section of HeLa Tom20-HaloTag cells stained with 1 µM JF585 for 30 min, fixed, embedded for EM and placed on a carbon-coated grid and fluorescence signals were registered by FM. Scale bars: A: 50 µm; B: 20 µm

**Fig. S5.**
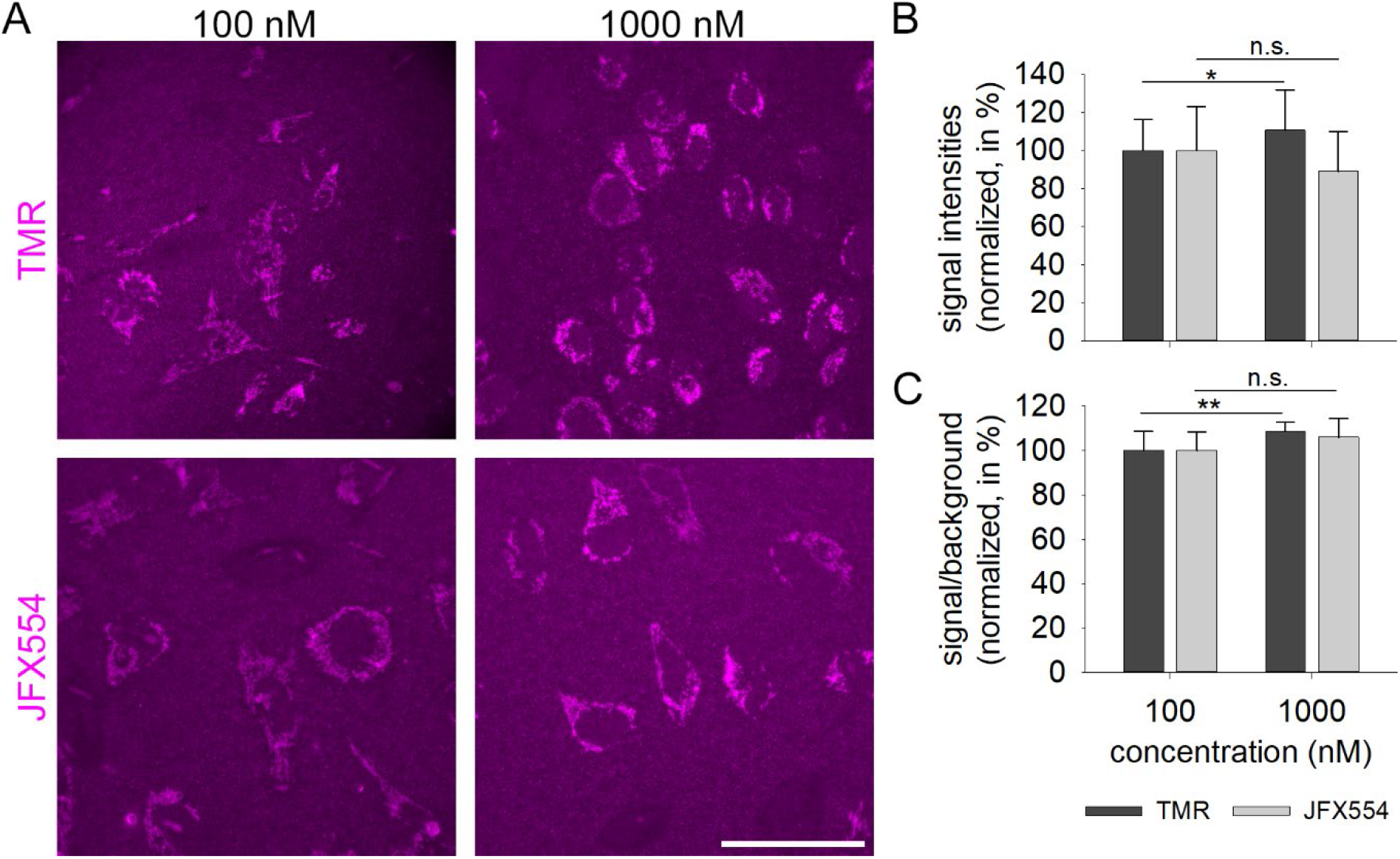
Increased dye concentrations improve in-resin fluorescence signals for TMR, but not for JFX554. HeLa Tom20-HaloTag cells stained with TMR or JFX554 in the indicated concentration for 30 min were further prepared as described for Fig. 1. **A**) Representative images of the respective dye and concentration. Scale bar, 50 µm. See Movie 5 for Z series. **B**) Signal intensities and **C**) signal-to-background ratio (S/B) were obtained as described for Fig. 1 and standardized by setting fluorescence signals obtained with 100 nM TMR or JFX554 to 100%. Statistical analyses were performed for each dye by unpaired, one-tailed t-test. Significances are indicated as follows: n.s., not significant, *, p < 0.05; **, p < 0.01; ***, p < 0.001.

**Fig. S6.**
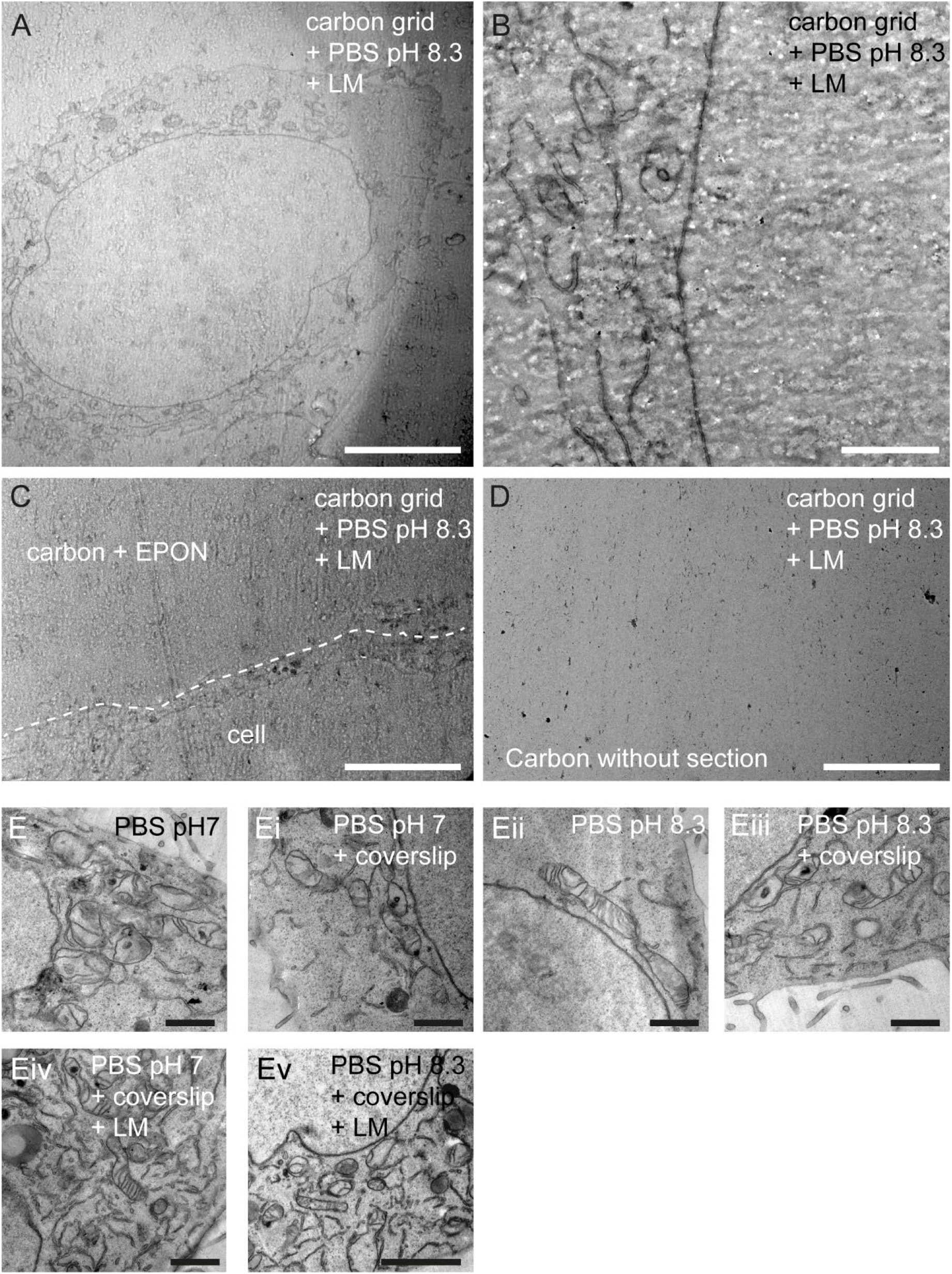
Artefacts induced after placing EPON sections on grids with carbon film only. HeLa cells were embedded as described in Material and Methods. Ultra-thin sections were prepared and mounted on commercial 200 mesh grids with carbon film (**A–D**), or on custom-made Formvar coated grids (**E, Ev**). Grids were subjected to the standard LM imaging (**A–D, Ev**), but also the influence of a different pH of the buffer (**E, Eii**), the use of the coverslip sandwich mentioned in Material and Methods (**Ei, Eii**), as well as the LM itself were tested on degradation of ultrastructure (**Eiv, Ev**). On grids with a carbon film, the EPON shows severe artefacts indicates by wavy or holey shapes (**A, B**). These artifacts were not restricted to cellular material, but appeared also in empty EPON (**C**), but not on the carbon film without a section (**D**). Different pH of the buffer or imaging conditions did not influence the ultrastructure on formvar coated grids (**E– Ev**). This hints to a general problem of the combination of EPON and carbon-coated grids. Scale bars: A, C, D: 5 µm; B, E, Ei, Eii, Eiii, Eiv: 1 µm; Ev: 2 µm.

**Fig. S7.**
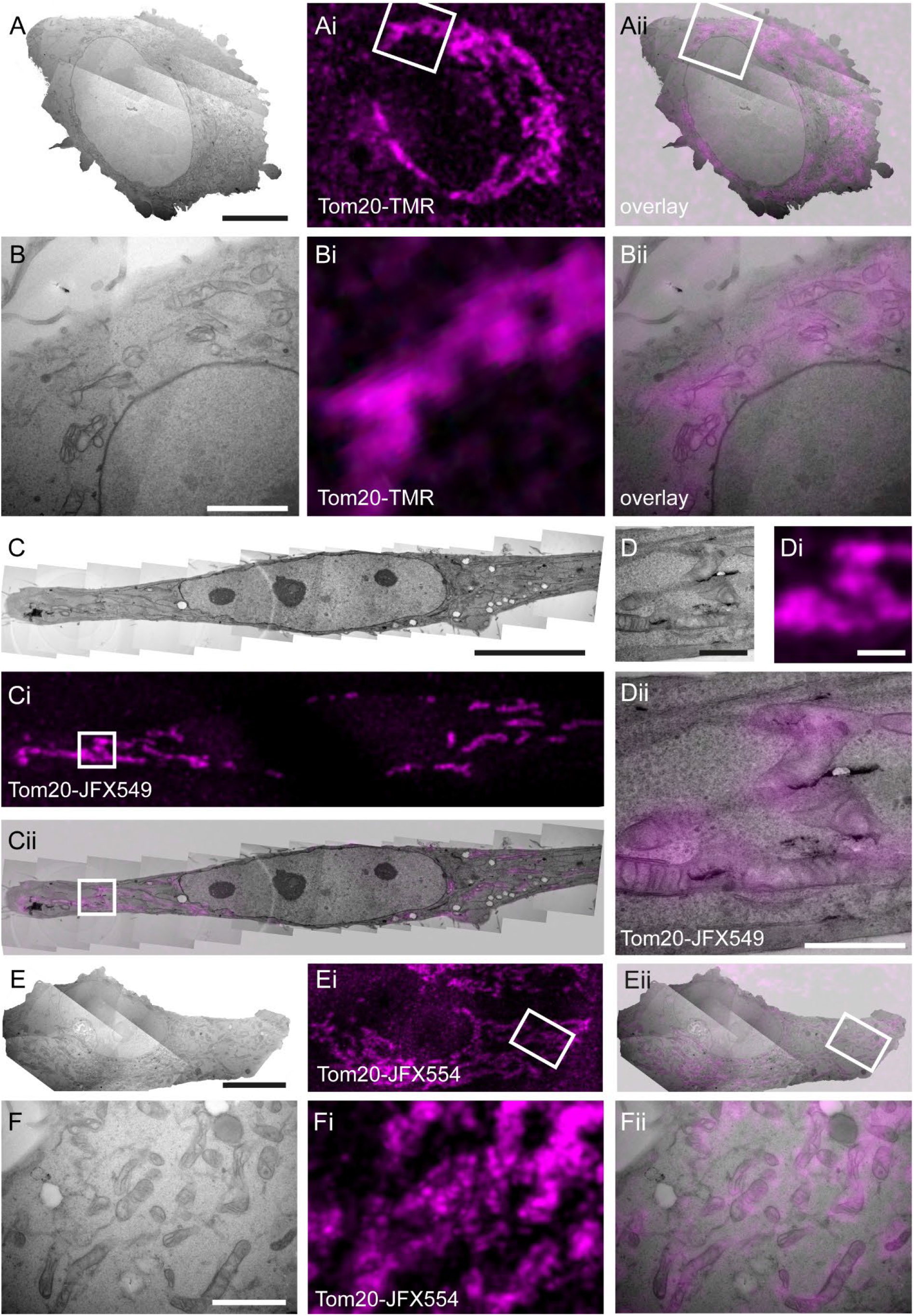
CLEM for Tom20-HaloTag using 250 nm sections with best-performing dyes and TMR. Hela Tom20-HaloTag cells were stained with 100 nM TMR (**A, B**), JFX549 (**C, D**) or JFX554 (**E, F**) for 30 min. After sample preparation for EM, 250 nm thin sections were prepared. Fluorescence signals were registered using CLSM. All three dyes retained their fluorescence after EPON embedding, and allowed correlation to mitochondria identified in TEM modality. Scale bars: A, E: 10 µm; B, F: 2 µm; C: 5 µm; D, Dii: 500 nm.

**Fig. S8.**
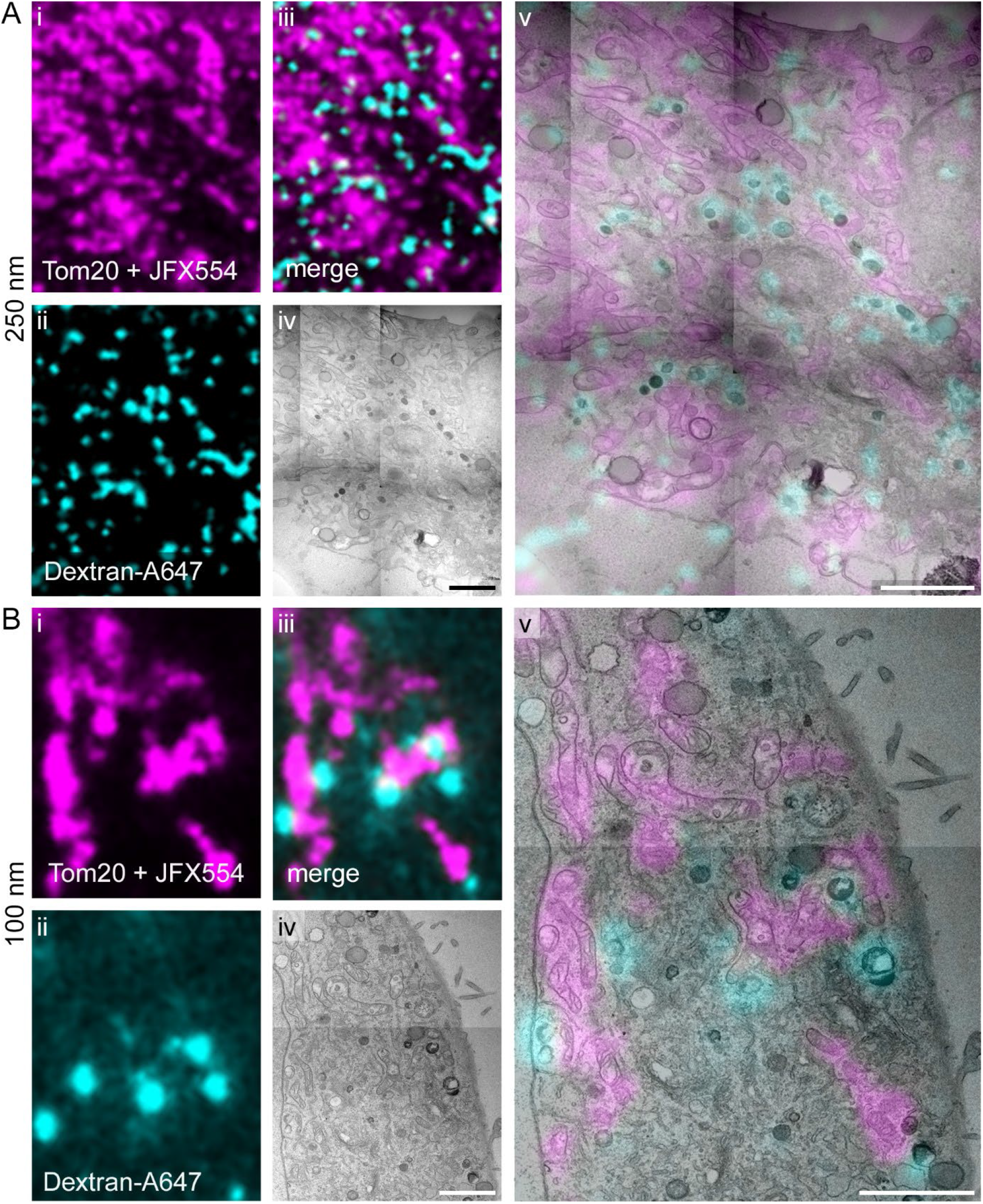
Dextran-Alexa Fluor 647 retains its fluorescence and is suitable for dual-color CLEM. Hela Tom20-HaloTag cells were pulsed with Dextran-Alexa Fluor 647 O/N and stained with 100 nM JFX554 for 30 min. Further sample preparation and imaging was conducted as described in Material and Methods. **A**) 250 nm section. **B**) 100 nm section. Dual-color CLEM and correlation of mitochondria and Alexa Fluor 647-containing endosomes was perfromed. Scale bars: 2 µm.

## Notes

### Competing Interest Statement

The authors have declared no competing interest.

